# How well do rudimentary plasticity rules predict adult visual object learning?

**DOI:** 10.1101/2022.12.31.522402

**Authors:** Michael J. Lee, James J. DiCarlo

## Abstract

A core problem in visual object learning is using a finite number of images of a new object to accurately identify that object in future, novel images. One longstanding, conceptual hypothesis asserts that this core problem is solved by adult brains through two connected mechanisms: 1) the re-representation of incoming retinal images as points in a fixed, multidimensional neural space, and 2) the optimization of linear decision boundaries in that space, via simple plasticity rules applied to a single downstream layer. Though this scheme is biologically plausible, the extent to which it explains learning behavior in humans has been unclear – in part because of a historical lack of image-computable models of the putative neural space, and in part because of a lack of measurements of human learning behaviors in difficult, naturalistic settings. Here, we addressed these gaps by 1) drawing from contemporary, image-computable models of the primate ventral visual stream to create a large set of testable learning models (n=2,408 models), and 2) using online psychophysics to measure human learning trajectories over a varied set of tasks involving novel 3D objects (n=371,000 trials), which we then used to develop (and [publicly release](http://www.github.com/himjl/hobj)) empirical benchmarks for comparing learning models to humans. We evaluated each learning model on these benchmarks, and found that learning models that using specific, high-level contemporary representations are surprisingly aligned with human behavior. While no tested model explained the entirety of replicable human behavior, these results establish that rudimentary plasticity rules, when combined with appropriate visual representations, have high explanatory power in predicting human behavior with respect to this core object learning problem.

**Author Summary:** A basic conceptual hypothesis for how an adult brain learns to visually identify a new object is: 1) it re-represents images as points in a fixed, multidimensional space, then 2) it learns linear decision boundaries that separate images of a new object from others, using a single layer of plasticity. This hypothesis is considered biologically plausible, but gauging its power to explain human learning behavior has not been straightforward. In part, this is because it is difficult to model how brains re-represent images during object learning. However, ongoing efforts in neuroscience have led to the identification of specific, image-computable models that are at least partially accurate descriptions of the neural representations involved in primate vision. Here, we asked whether any of those representations, when combined them with simple plasticity rules, could make accurate predictions over a large body of human object learning behavioral measurements. We found that specific models could indeed explain a majority of our behavioral measurements, suggesting the rudimentary, biologically-plausible mechanisms considered here may be sufficient to explain a core aspect of human object learning.

## Introduction

People readily learn to recognize new visual objects. As an individual receives views of a new object – new spatial patterns of photons striking their eyes – their ability to correctly recognize new views of that object increases, possibly very rapidly. What are the mechanisms that allow an adult human to do so?

Efforts from cognitive science, neuroscience, and machine learning have led to a diverse array of ideas to understand and replicate this human ability, and human example-based learning in general. These works range in levels of specification, from conceptual frameworks that do not directly offer quantitative predictions [1–5], models which depend on unspecified intermediate computations (i.e. non-image-computable models; [6–10]), to end-to-end learning models which take images as input [11–17].

An important step in determining the extent to which any of these ideas are valid descriptors of human object learning (and its underlying neural mechanisms) is to evaluate them on the basis of their ability to predict empirical measurements of human behavior, over a range of task settings. While such model evaluations (which we refer to as “benchmarks”) exist for visual tasks involving known object categories (e.g. [18–20]), the field currently lacks a readily available set of benchmarks for visual tasks involving the learning of novel objects.

To begin addressing this gap, we designed novel benchmarks for human object learning. We focused on object learning in the context of binary discrimination tasks, in which a learner must acquire the ability to discriminate two new objects from each other, receiving feedback (correct or incorrect) on each trial. Though this elementary task paradigm does not encompass all possible aspects of “object learning” per se, it exposes subjects to one of its core problems: using a finite number of images of a new object to accurately identify that object in novel images.

Our first experimental goal was to measure this ability in adult humans across several tasks, each involving highly varied views of novel 3D objects. The resultant set of measured learning curves (one for each task) formed a large set of human “behavioral signatures” which could then be used as the basis for our first and primary benchmark. Based on the extent a candidate learning model quantitatively reproduced (or failed to reproduce) those human signatures, its empirical validity could be established.

Next, motivated by prior suggestions of where humans may be particularly powerful [13, 21], our secondary experimental goal was to take additional measurements of human behavior during the special case of “one-shot” learning, where the subject receives just a single view (that is, a single image) of each object before being asked to generalize to unseen views of that object. Here, we sought to measure human generalization over a variety of tests involving identity-preserving transformations (e.g. translation, scaling, and 3D rotation) in this one-shot setting, then to use the resultant measurements to create our secondary benchmark, which evaluates a model’s ability to replicate human-like patterns of generalization across the same tests.

Once we generated these two benchmarks, we used them to evaluate a longstanding conceptual hypothesis for object learning, which posits that adult humans re-represent incoming visual stimuli in a stable, multidimensional Euclidean space, learn linear categorization boundaries in that space, and apply this learned boundary to categorize new stimuli [5, 7, 22].

This overall framework is notable in that it has plausible neural implementations. For example, the re-representation of an image as a location in a multidimensional space could be expressed by the pattern of population firing rates in a visually-driven area; linear categorization boundaries could be implemented by downstream neurons that respond to weighted sums of upstream activity [23]; and those weights could be adjusted through simple, reward-driven plasticity mechanisms [24–26].

However, despite the biological plausibility of this framework, it does not directly generate quantifiable predictions of human behavior. To generate such predictions, the representational space thought to be used by humans must first be specified, as well as the precise form of the plasticity rule. While there are standard learning algorithms that can be written as reward-based plasticity rules operating on a single layer (e.g. [27]), obtaining an accurate specification of the representational space presents a larger challenge, in that this space may have a highly complex relationship with the content of images.

Typically, this challenge has been approached by limiting the domain of the learning model to a restricted, often simple domain of images, then assuming that the axes of the representational space correspond to latent variables used to generate those specific images (e.g. [10]). Alternatively, data-driven approaches have been used to estimate the embedding of those specific images in the representational space (see [3]). However, it has not been clear how to extend those approaches to the domain of all arbitrary images, including the kinds of images considered here, or whether any learning models based on the proposed mechanisms would offer accurate predictions of human behavior in more complex and naturalistic visual domains.

To address this gap, we drew from ongoing efforts in cognitive science [28–33] and visual neuroscience [34–36] that have established correspondences between the intermediate layers of image-computable, deep convolutional neural networks (DCNNs) and human visual behavior, as well as the activity of neural populations across the primate ventral stream.

We took a large sample of those image-computable layers and combined each of them with a variety of simple plasticity rules (drawn from statistical learning theory and reinforcement learning) to create a battery of testable learning models. If humans indeed use a representation that is sufficiently well-approximated by one of those intermediate layers and have plasticity mechanisms that are well-described by one of these plasticity rules, then the corresponding learning model should have close empirical correspondence with observed human behaviors, as evaluated by the benchmarks here.

At the outset of this study, we reasoned that, if the behavior produced by any model was found to be indistinguishable from that of humans, it could serve as a leading scientific hypothesis to drive further experiments. If they were not found, predictive gaps could be used to guide future work in improving models of human object learning. Either way, the benchmarks created in this work could facilitate a standard evaluation of current and future visual object learning models.

## Results

We measured human behavior over two variants of an object learning task (Experiments 1 & 2). In Experiment 1, we measured human subjects learning to discriminate pairs of novel objects as subjects were provided with an increasing number of views of those objects, and feedback on their choices. In Experiment 2, we also measured humans learning to discriminate between pairs of objects, but provided only one view per object before assessing subjects’ accuracy on a variety of generalization tests. The results of each experiment are presented below, along with quantitative comparisons of those results with a large set of learning models. Further details of the experiments and the models are provided in the Materials and methods.

### 0.1 Experiment 1: Humans are rapid, but imperfect novel object learners

In Experiment 1, we measured a population of anonymous human subjects (n=70) performing 64 learning subtasks. Each subtask required that the subject learn to discriminate a different pair of two novel objects, rendered under high view variation (see Fig 1A).

**Fig 1.**
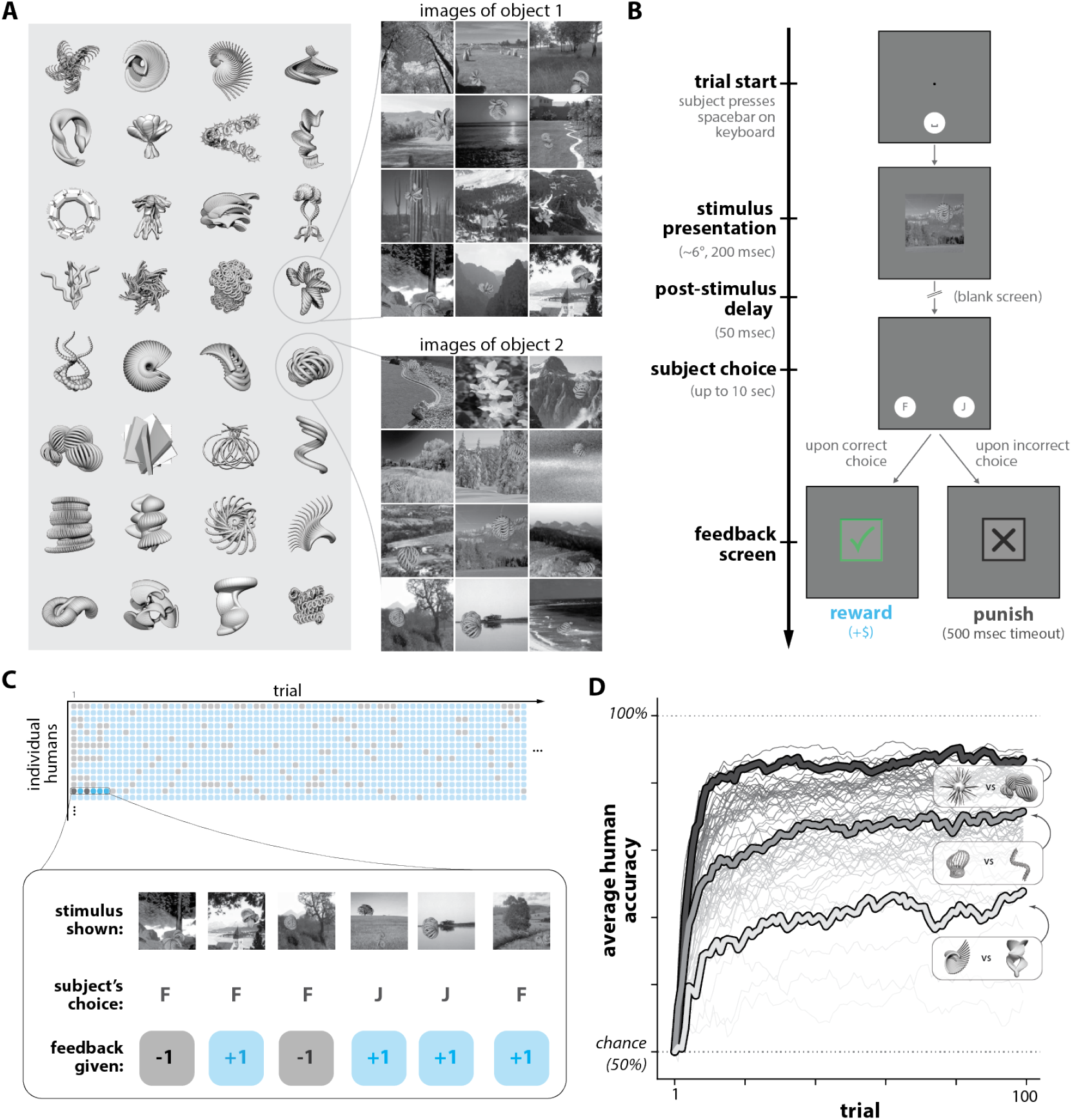
Humans learning novel objects. **A. Images of novel objects**. Images of synthetic 3D object models were created using random viewing parameters (background, location, scale, and rotational pose). **B. Task paradigm**. On each trial, a randomly selected image (of one of two possible objects) was briefly shown to the subject. The subject had to report the identity of the object by making one of two possible choices (“F” or “J”). Positive reinforcement was delivered if the subject choice was “correct”, per an object-choice contingency that the subject learned through trial-and-error (e.g., object 1 corresponds to “F”, and object 2 corresponds to “J”). **C. Example subject-level learning data**. Each subject performing a subtask performed a randomly sampled sequence of 100 trials (i.e. images and their choice-reward contingencies), and we measured their sequence of correct (blue) and incorrect (gray) choices. Image stimuli were never repeated, ensuring each trial tests the subject’s ability to generalize to unseen views of the objects. **D. Human learning curves**. We averaged across human subjects to estimate accuracy as a function of trials for n=64 subtasks (each consisting of a distinct pair of objects). We found that some subtasks were found to be reliably harder for humans than others; three example subtasks across the range of difficulty are highlighted. Learning curves shown are smoothed with a moving window filter (for visualization only).

These subtasks proceeded in a trial-by-trial fashion. At the beginning of a trial, a test image containing one of two possible objects was briefly presented (at ≈6°of the visual field for ≈200 milliseconds). Then, the subject was asked to report which of the two objects was present “in” that image through a button press, and evaluative feedback (correct or incorrect) was delivered based on their choice (see Fig 1B). The subtask then proceeded to the next trial (for a total of 100 trials).

The core measurement we sought to obtain for each subtask was the discrimination accuracy of a typical subject as a function of the number of previously performed trials (i.e. the *learning curve* for each subtask). To estimate the learning curve for a particular subtask, we recorded the sequence of corrects and incorrects achieved by achieved by multiple subjects (n=50 subjects) performing n=100 randomly sampled trials (see Fig 1C), then averaged across subjects to estimate the learning curve for that subtask. We estimated learning curves in this manner for all n=64 subtasks in this experiment (depicted in Fig 1D).

Upon examination of these learning curves, we found that on average (over subjects and subtasks), human discrimination accuracy improved immediately – i.e. after a single image example and accompanying positive or negative feedback. By construction, accuracy on the first trial is expected to be 50% (random guessing); but on the following trial, humans had above-chance accuracy (mean 0.65; [0.63, 0.67] 95% bootstrapped CI), indicating behavioral adaptation occurred immediately and rapidly. Average discrimination accuracy continued to rise across learning: the subject-averaged, subtask-averaged accuracy on the last trial (trial 100) was 0.87 (mean; [0.85, 0.88] 95% CI). The subject-averaged, subtask-averaged accuracy over all 100 trials was 0.82 (mean; [0.81, 0.84] 95% CI).

As anticipated, we found that *different* subtasks (i.e. different pairs of objects) could have widely different learning curves. This is illustrated in Fig 1D, which shows the estimated average human learning curve for each subtask. That is, we observed that some tasks were “easy” for humans to learn, and some were harder (e.g. mean accuracy of ≈0.65 for the most difficult 10% of subtasks). These variations were not artifacts of experimental variability, which we established by estimating the value of Spearman’s rank correlation coefficient between average subtask performances that would be expected upon repetitions of the experiment (*ρ* = 0.97; see S2 Appendix). Moreover, we found our measures of human learning behavior were robust to the experimental imprecision in online behavioral testing (e.g. from head movement; variation in monitor setups) by replicating a subset of this experiment in a population of in-lab subjects with head-fixation and eye tracking (see S1 Fig).

Overall, these observations indicate that 1) humans can acquire a significant amount of learning with respect to novel visual object concepts with a small number of examples (e.g. 4 training examples to reach 75% correct, 6 to reach 90% of their final performance), and 2) learning new objects is highly dependent on the 3D shapes of those objects, with many object pairs being far from perfectly learned within 100 trials.

We next asked how well a family of models based on a standard cognitive theory of learning are – or are not – able to explain these behavioral measurements.

### 0.2 Comparing image-computable learning models with humans

As described above, the core set of behavioral measurements we obtained in Experiment 1 consisted of subject-averaged learning curves for each subtask (accuracy values for 64 subtasks across 100 trials). Our next step was to to specify a procedure that assesses how well any given computable model of object learning can quantitatively reproduce those curves.

Given a model, this procedure consisted of 1) simulating the same set of subtasks in the model and estimating its learning curves for those subtasks (exactly analogous to how they were estimated in humans), and 2) computing a scalar-valued error score (MSE_*n*_) which summarizes the extent to which the learning curves of the model quantitatively matches (or not) the corresponding learning curves in humans.

We sought to score models drawn from a standard model family (see Fig 2A). Each model in this family consists of two conceptual stages: which we refer to as 1) an *encoding stage* which places each incoming image into a representational space, and 2) a tunable *decision stage*, which generates a choice by using a linear decision boundary in that space. The learning of new object-choice associations is guided by a *plasticity rule*, which processes environmental feedback (the same feedback information provided to human subjects) to adjust the linear weights of the tunable decision stage. Plasticity occurs only in the decision stage; the encoding stage is held completely fixed.

**Fig 2.**
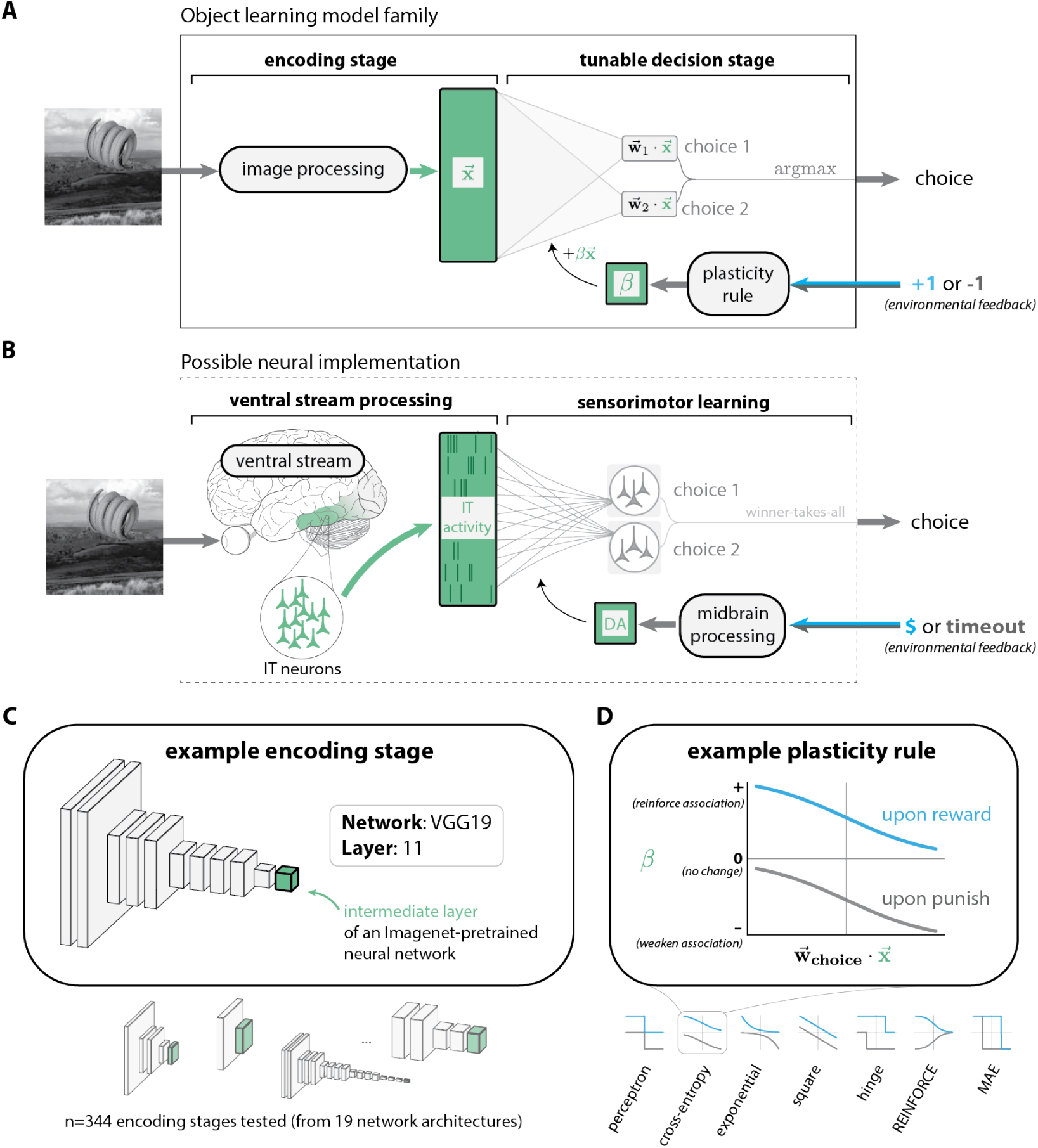
Model family of object learning in humans. **A. Model family**. Each model in this family had two stages: an *encoding stage* which re-represents an incoming pixel image as a vector 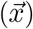, and a tunable *decision stage*, which uses 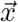 to generate a choice by computing linear choice preferences 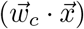, then selecting the most preferred choice. Subsequent environmental feedback is processed to update the parameters of the decision stage. Specific models in this family correspond to specific choices of the encoding stage and plasticity rule. **B. Possible neural implementation**. The functionality of the encoding stage in **A** could be implemented by the ventral stream, which re-represents incoming retinal images into a pattern of distributed population activity in high level visual areas, such as area IT. Sensorimotor learning might be mediated by plasticity in a downstream association region. Midbrain processing of environmental feedback signals could guide plasticity via dopaminergic (DA) projections. **C. Encoding stages**. We built learning models based on several encoding stages, each based on a specific intermediate layer of an Imagenet-pretrained deep convolutional neural network. **D. Plasticity rules**. We drew basic plasticity rules from statistical learning theory and reinforcement learning (Table 0.13.2). Each rule aims to achieve a slightly different optimization objective (e.g., the “square” rule attempts to minimize the squared error between the choice preference and the subsequent magnitude of reward).

**Fig 3.**
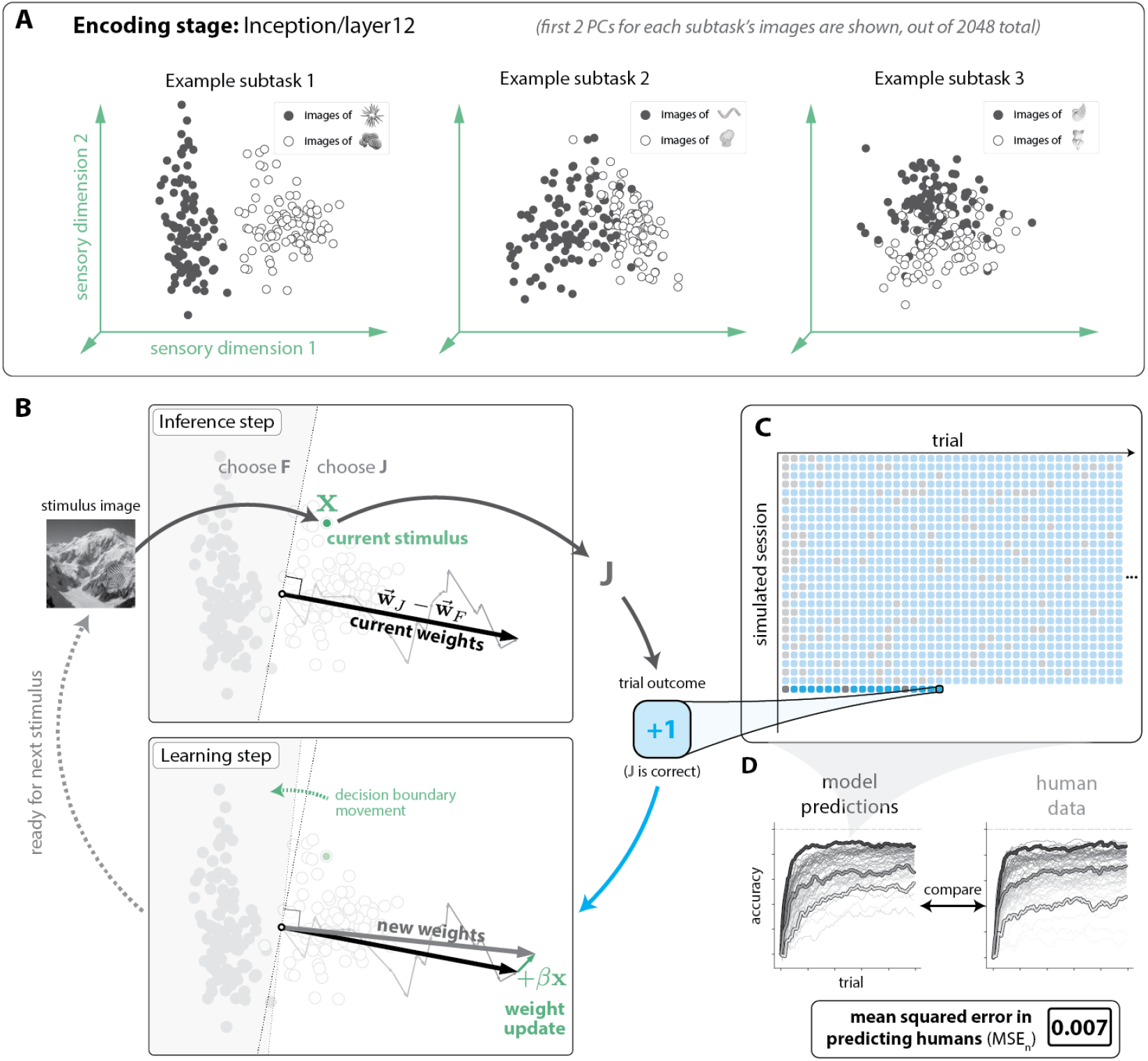
Model simulations of human learning. **A. Example encoding stage representations of novel object images**. Each subtask consists of images of two novel objects (indicated in black and white dots). The first two principal components of a 2048-dimensional encoding stage (Inception/layer12) are shown here (computed separately on the images for each subtask, for clarity). Linear separability can be observed, to varying degrees. **B. Simulating a single trial**. Clockwise, starting from top left: the incoming stimulus image is re-represented by the encoding stage into a location in a representational space, 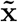. Preferences for each choice are computed (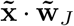 and 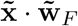), and the most preferred choice is selected, which amounts to making the choice based on where 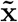 falls with respect to a linear decision boundary 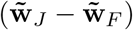. Here, the choice “J” is selected, which happens to be the correct choice for this image. The subsequent reward causes the decision boundary to change based on the plasticity rule. **C. Simulated model behavioral data**. For each learning model, we simulated a total of n=32,000 behavioral sessions (64 subtasks, 500 simulations each), and recorded its behavior (correct or incorrect) on each trial. **D. Comparing model and human behavior**. We averaged across simulations to obtain the model’s learning curves for each subtask, then compared them to subject-averaged human learning curves, using a bias-corrected mean-squared error metric (MSE_*n*_; see Bias-corrected mean squared error) to quantify the (dis)similarity of the model to humans.

Here, we implemented learning models based on different combinations of encoding stage and plasticity rule. We considered n=344 encoding stages based on specific intermediate layers of Imagenet-pretrained DCNNs [37], and n=7 plasticity rules drawn from statistical learning theory and reinforcement learning (see Baseline model family for details). In total, we implemented models based on all possible combinations of these encoding stages and plasticity rules (n=2,408 learning models).

We then tested each model on the same set of subtasks as humans, simulating n=32,000 behavioral sessions per model (n=500 simulations per subtask). We estimated each model’s learning curves for each subtask (i.e. accuracy values over 64 subtasks and 100 trials), and scored its similarity to humans using a mean-squared-error statistic (MSE_*n*_; details in Bias-corrected mean squared error). We show an illustration of this procedure for an example model shown in Box 3, and a histogram of MSE_*n*_ scores for all tested models is shown in Fig 4.

**Fig 4.**
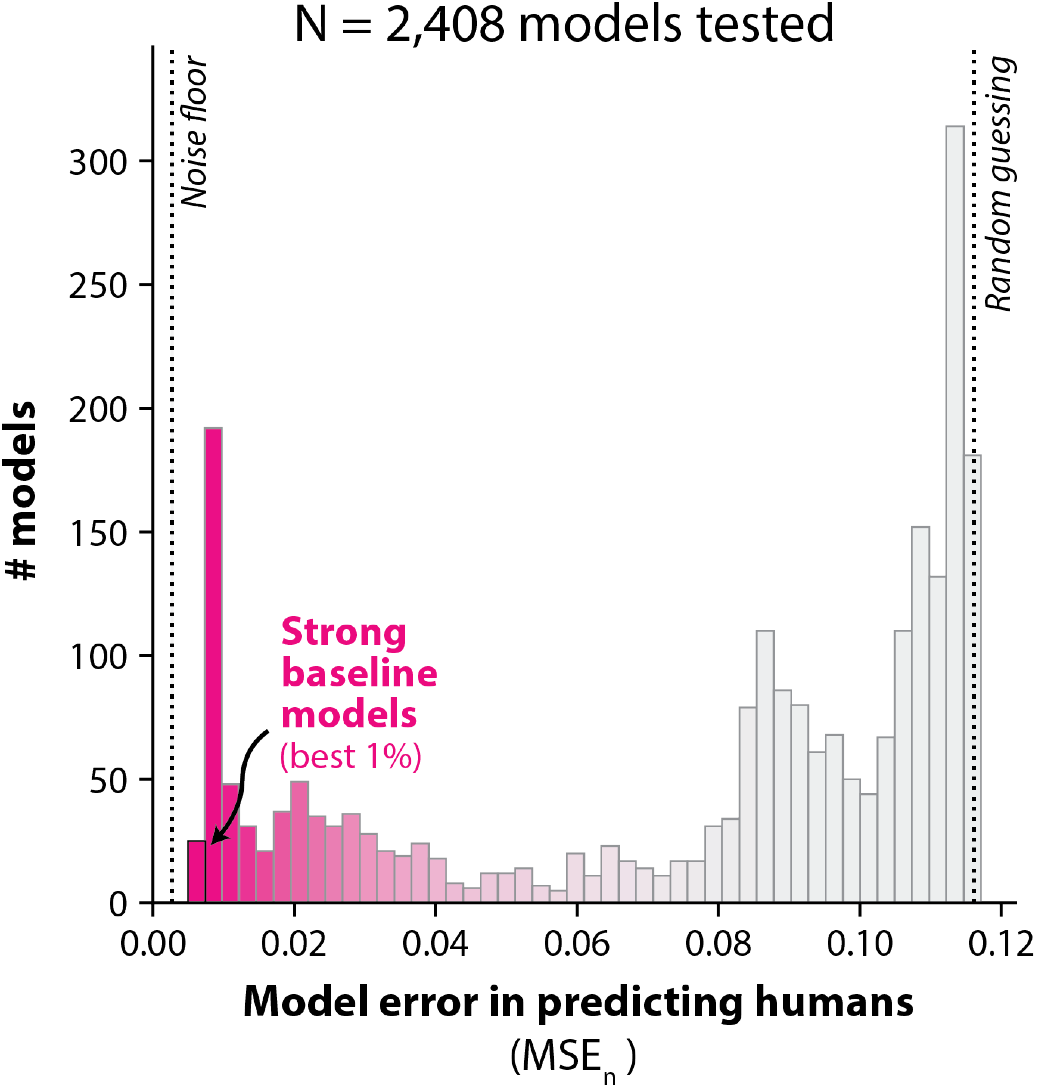
Bias-corrected mean-squared errors (MSE_*n*_) vs. humans for all models tested. The n=2,408 models we tested varied widely in the extent of their alignment with human learning (i.e. their average squared prediction error). We denote the best 1% of such models as *“strong baseline models”*. The noise floor corresponds to an estimate of the lowest possible error achievable 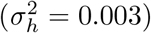, given the experimental power in this study. The vertical line labeled “random guessing” marks the error incurred by a model which produces a random behavioral output on each trial.

A higher value of MSE_*n*_ means the model is a worse predictor of human learning behavior; lower values are better. In principle, no model can be expected to have an MSE_*n*_ lower than a “noise floor”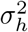, which comes from the uncertainty in our experimental estimates of each human learning curve (i.e. from the finite amount of human data collected). Intuitively, the value of 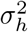 can be understood as the MSE_*n*_ score that can be expected from the “perfect” model of human learning behavior. We made an unbiased estimate of this noise floor (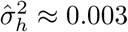, see Noise floor estimation), then compared this to the MSE_*n*_ scores achieved by the models we tested. We note that the *root* value of the noise floor (and MSE_*n*_) is a rough estimate of the expected error in the native units of the measurements (accuracy values).

Many of the models were far from the noise floor, but we found that a subset of models achieved relatively low error. For example, we found the best 1% of the models (which we refer to as “strong baseline models”) had root-mean squared errors of 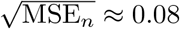, coming relatively close to the noise floor 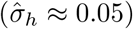. Still, all models, including these ones, were statistically distinguishable from humans; all models were rejected as having expected behavior identical to humans with significance level of at least *p* < 0.001 (see Null hypothesis testing).

#### 0.2.1 Model components affecting the score of a model

Given the range of MSE_*n*_ scores we observed, we next wished to perform a secondary analysis on how each of the two components defining a model (its encoding stage and plasticity rule) affected its error score on the benchmark above. One general trend we observed was that models built with encoding stages from deeper layers of DCNNs tended to produce more human-like learning behavior (see Fig 2B). On the other hand, the choice of plasticity rule (which defined by the tunable decision stage) appeared to have little effect in a model’s ability to generate human-like learning behavior (see Fig 5A).

**Fig 5.**
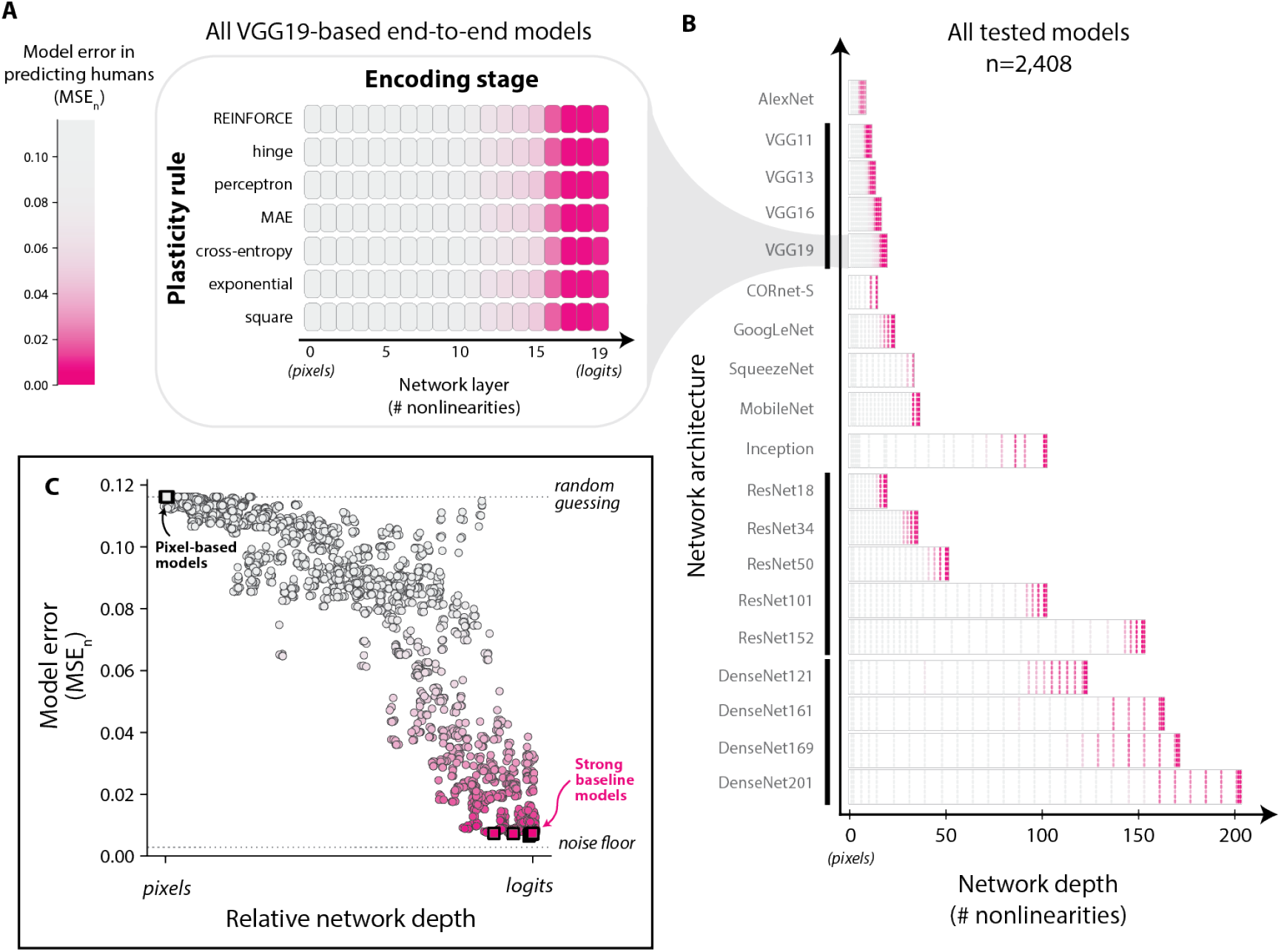
Evaluating the effect of model design choices on predictive accuracy of human learning. **A. Example model scores across encoding stages and plasticity rules**. A typical example of the relative contributions of plasticity rule (y-axis) and encoding stage (x-axis) on model scores (MSE_*n*_, encoded by color). **B. Overview of all models tested**. In total, we tested encoding stages drawn from a total of n=19 DCNN architectures, varying widely in depth. A model’s similarity to humans was highly affected by the choice of encoding stage; those based on deeper layers of DCNNs showed the most human-like learning behavior. On the other hand, the choice of plasticity rule had a minuscule effect. **C. Predictive accuracy increases as a function of relative network depth**. Learning models with encoding stages based on DCNN layers closer to the final layer of the architecture tended to be better.

We quantified these observations by performing a two-way ANOVA over all model scores (see S1 Appendix), treating the plasticity rule and encoding stage as the two factors. This analysis showed that the choice of plasticity rules explained less than 0.1% of the variation in model scores; by contrast, 99.8% of the variation was driven by the encoding stage, showing that the predominant factor defining the behavior of the learning model was the encoding stage.

#### 0.2.2 Strong baseline models are largely, but not perfectly, correlated with human performance patterns

The set of MSE_*n*_ scores indicated that *all* models tested had significant differences from humans, based on their respective learning curves. There are several ways in which these differences could originate – for example, a model might have “fast” learning curves for tasks that humans learn slowly, and “slow” learning curves for tasks that humans learn rapidly. Alternatively, a model might have the same pattern of difficulty across subtasks as humans, but simply be slower at learning, overall.

To gain insight into these possibilities, we performed an additional analysis in which we compared each model to humans along two more granular statistics: 1) its *overall accuracy* over all subtasks and trials tested, and 2) its *consistency* with the patterns of difficulty exhibited by humans across subtasks (see Fig 6A).

**Fig 6.**
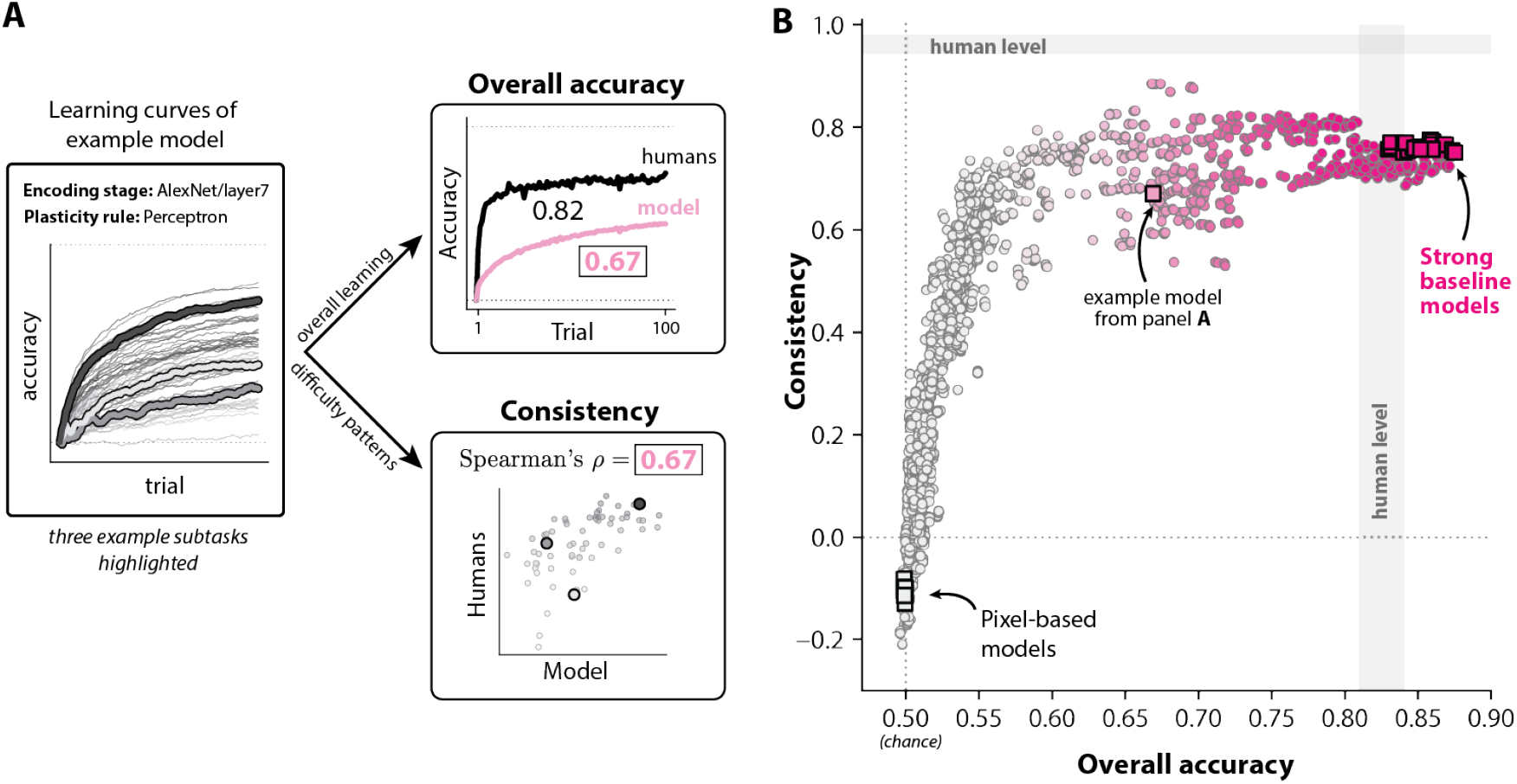
Comparing models to humans along more granular behavioral signatures of learning. **A. Decomposing model behavior into two metrics**. We examined model behavior along two specific aspects of learning behavior: **overall accuracy** (top right), which is the average accuracy of the model over the entire experiment (i.e. averaging across all 100 trials and all 64 subtasks), and **consistency** (bottom right), which conveys how well a model’s pattern of trial-averaged performance over different subtasks rank-correlates with that of humans (Spearman’s rank correlation coefficient). **B. Consistency and overall accuracy for all models**. Strong baseline models (top-right) matched (or exceeded) humans in terms of overall accuracy, and had similar (but not identical) patterns of performance with humans (consistency). The gray regions are the bootstrap-estimated 95% range of each statistic between independent repetitions of the behavioral experiment. The color map encodes the overall score (MSE_*n*_) of each model (colorbar in Fig 5A).

Intuitively, overall accuracy is a gross measure of a learning system’s overall ability to learn, ranging from 0.5 (chance, no learning occurs) to 0.995 (learning completed after just one trial). Consistency (*ρ*) quantifies the extent to which a model finds the same subtasks easy and hard as humans, and ranges from *ρ* = −1 (perfectly anticorrelated pattern of performance) to *ρ* = −1 (perfectly correlated pattern of performance). A value of *ρ* = 0 indicates no correlation between the patterns of difficulty across subtasks in a model and humans.

These metrics are theoretically unrelated to each other; given any overall accuracy,^1^ a model may have a high or low consistency with humans, and vice versa. Nevertheless, we observed that these two metrics strongly covaried for these models; models with high overall accuracy also tended to have high consistency (see Fig 6B).

#### 0.2.3 Humans learn new objects faster than all tested models in low-sample regimes

Though many models matched or exceeded human-level overall accuracy (i.e. accuracy averaged over trials 1-100 for all subtasks), we noticed that all models’ accuracy *early on* in learning consistenctly appeared to be below that of humans (see Fig 7A). We tested for this by comparing the accuracy of models and humans in an initial phase of learning (trials 1-5 for all subtasks), and indeed found that *all* of the models were significantly worse than humans in the early phase of learning (all p<0.05, bootstrap hypothesis test, see Fig 7B).

**Fig 7.**
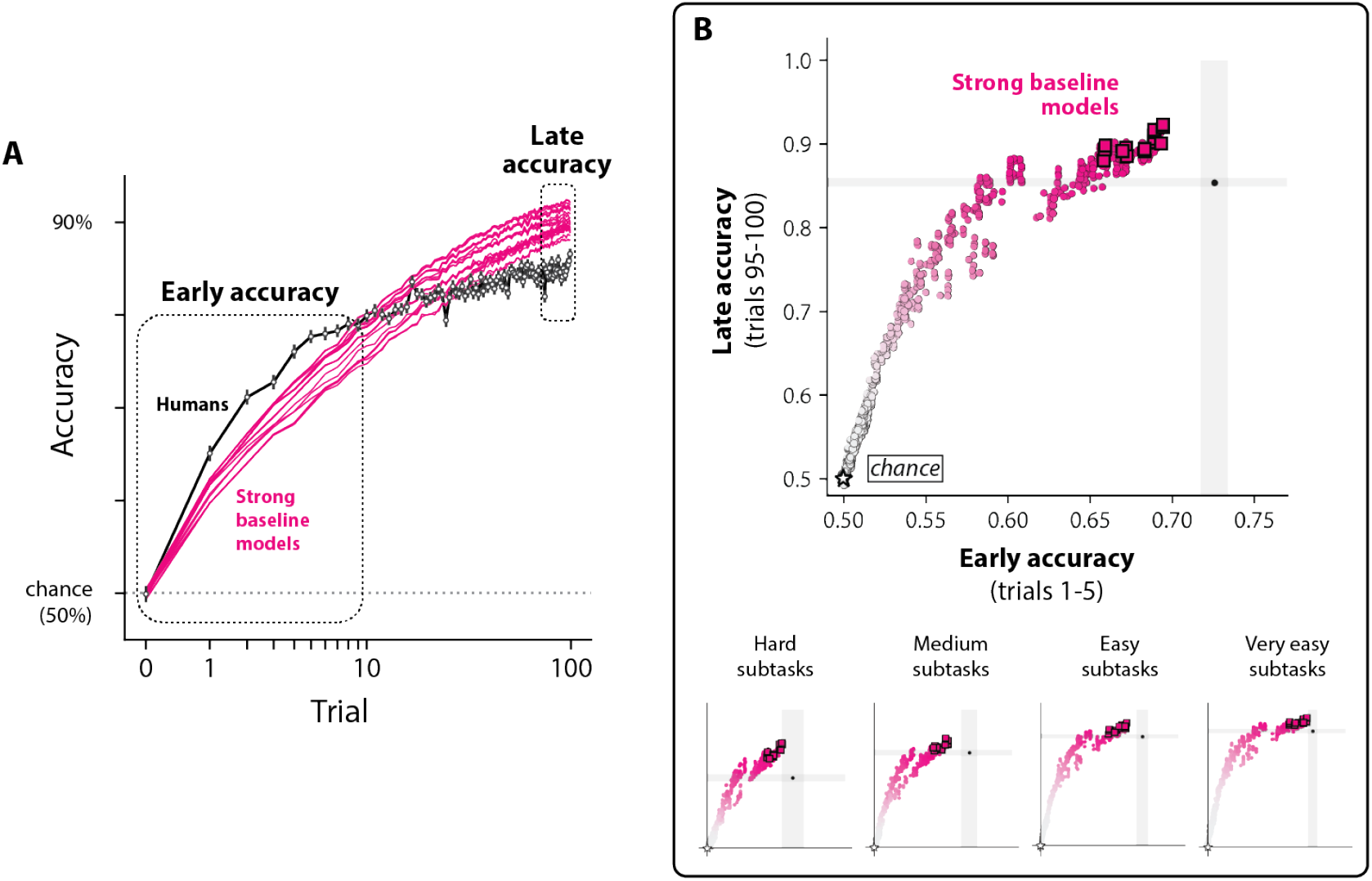
Humans outperform all strong baseline models in low-sample regimes. **A. Subtask-averaged learning curves for humans and strong baseline models**. The y-axis is the percent chance that the subject made the correct object report (chance is 50%). The x-axis is the total number of image examples shown prior to the test trial (log scale). The average learning curves for humans (black) and models (magenta) show that humans outperform all models in low-sample learning regimes. Errorbars on the learning curves are the bootstrapped SEM; model errorbars are not visible. **B. No model achieves human-level early accuracy**. We tested all models for whether they could match humans early on in learning. Several models (including all strong baseline models) were capable of matching or exceeding late accuracy in humans (average accuracy over trials 95-100), but no model reached human-level accuracy in the early regime (average over trials 1-5). This trend was present in subtasks across different levels of difficulty (bottom row). The gray region shows the 95% bootstrapped CI for each statistic; 95% CIs for models are too small to be shown.

We wondered whether this gap was present across levels of difficulty (e.g., that models tended to perform particularly poorly on “hard” subtasks relative to humans, but were human-level for other subtasks), and repeated this analysis across four different difficulty levels of subtasks (where each level consisted of 16 out of the 64 total subtasks we tested, grouped by human difficulty levels). We found models were consistently slower than humans across the difficulty range, though we could not reject a subset (11/20) of the strong baseline models at the easiest and hardest levels (see Fig 7B).

Lastly, though all models failed to match humans in the early regime, many models readily matched or exceeded human performance late in learning (i.e. the average accuracy on trials 95-100 of the experiment).

### 0.3 Experiment 2: Characterizing one-shot object learning in humans

Our observation above suggested these models learn more slowly than humans in few-shot learning regimes involving random views of novel objects. To further characterize possible differences between models and humans in this “early learning” regime, we performed an additional behavioral experiment (Experiment 2) in which we measured the ability of humans to generalize following experience with only a single image of each object category.

Experiment 2 followed the same task paradigm as Experiment 1 (binary discrimination learning with evaluative feedback). Each behavioral session was based on one of 32 possible subtasks (i.e. 32 possible pairs of novel objects), and began with a “training phase” of 10 trials in which the subject acquired an object-response contingency using a single canonical image for each of the two objects. After the training phase, we then asked subjects to perform trials with “test images” consisting of transformed versions of the two training images (see One-shot behavioral testing for further details).

These test images were generated by applying the five kinds of image variations present in our original experiment (translation, scale, random backgrounds, in-plane object rotation, and out-of-plane object rotation) to the test images. We also generated test images using four additional kinds of image variation that were not present in the original experiment (contrast shifts, pixel deletion, blur, and shot noise), but might nonetheless serve as informative comparisons for identifying functional deficiencies in a model relative to humans. For each kind of transformation, we tested four “levels” of variation. For example, in measuring humans’ one-shot test accuracy to scale, we showed subjects images where the object was resized to 12.5%, 25%, 50%, and 150% of the size of the object in the original training image (see One-shot stimulus image generation for details).

Under this experimental setup, the core measurements we sought to obtain was the subject-averaged discrimination accuracy for n=36 generalization tests (9 transformation types, 4 levels of variation each). Intuitively, each of the n=36 measurements is an estimate of a typical human’s ability to successfully generalize to a specific kind and magnitude of view transformation (e.g. downscaling the size of an object by 50%), after exposure to a single positive and negative example of a new object. Unlike Experiment 1, here we combined observations across the 32 subtasks used in this experiment, ignoring the fact that there may be variation in these measurements based on the specific objects involved. We also attempted to correct for any memory or attentional lapses in these estimates (see One-shot behavioral statistics in humans for details).

Across these 36 test conditions, we found that humans had varied patterns of generalization (Fig 8B). For example, we observed that accuracy varied systematically based on the level of variation applied with respect to scale, out-of-plane rotation, and blur. On the other hand, human subjects had nearly perfect generalization across all tested levels of variation for translations, backgrounds, contrast shifts, and in-plane rotations. Overall, these diverse patterns of generalization were estimated with a relatively high degree of experimental precision, as quantified by our estimates of the human noise floor for this experiment (root noise floor of 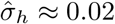).

**Fig 8.**
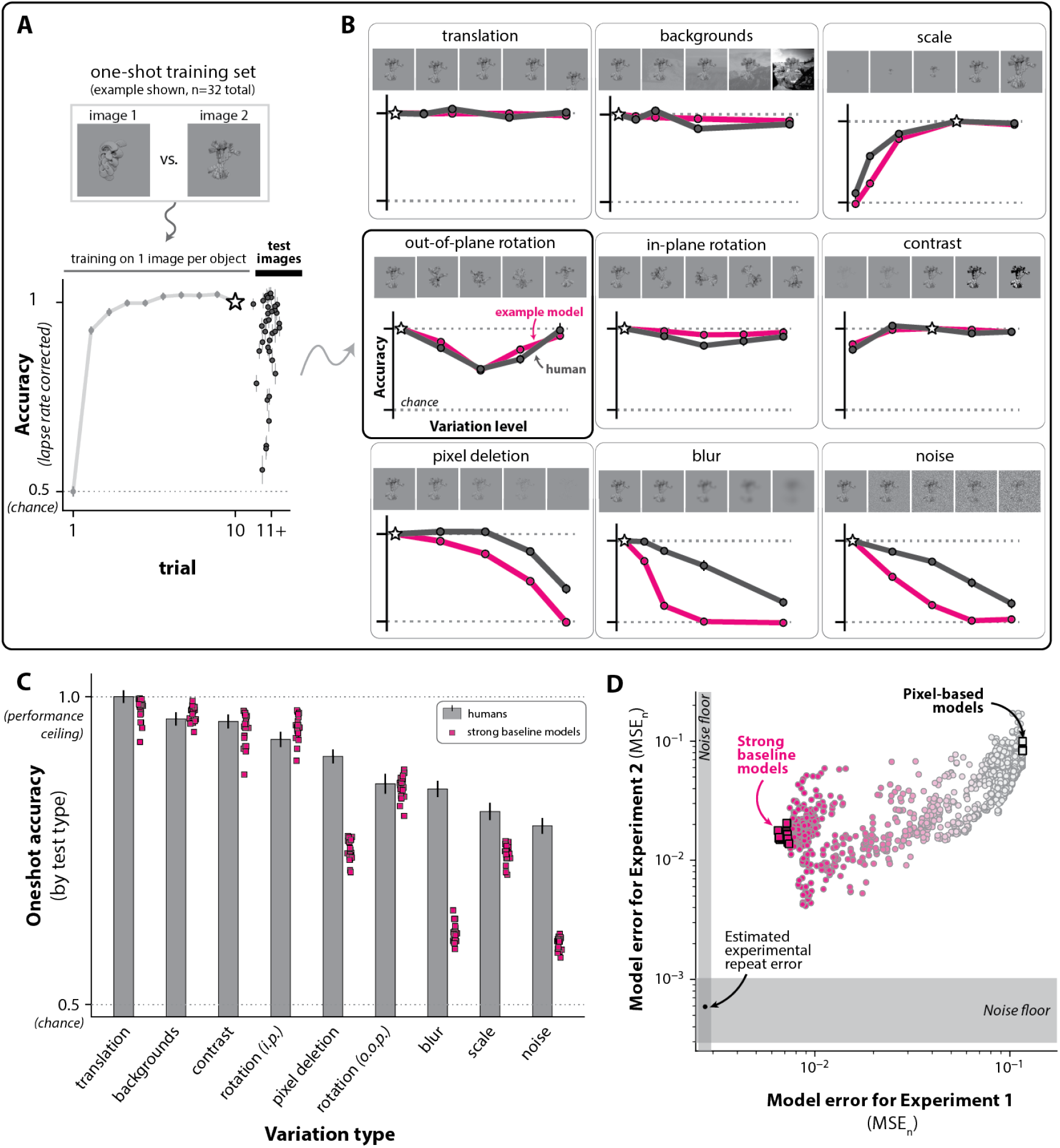
One-shot learning in humans. **A. One-shot learning task paradigm**. We performed an additional study (Experiment 2) to characterize human one-shot learning abilities (using the same task paradigm in Fig 1). The first 10 trials were based on two images (n=1 image per object) that were resampled in a random order. On trials 11-20, humans were tested on transformed versions of those two images (nine types of variation, four variation levels, n=36 total generalization tests) **B. Human and example model one-shot accuracy for all generalization tests**. An example strong baseline model’s pattern of generalization (magenta) is shown overlaid against that of humans. **C. Humans outperform strong baseline models on some kinds of image variations**. We averaged human one-shot accuracy (gray) on each type of image variation, and overlaid all strong baseline models (magenta). The errorbars are the the 95% CI (basic bootstrap). **D. Comparison of MSE**_*n*_ **scores for Experiment 1 and 2**. No strong baseline model could fully explain the pattern of one-shot generalization observed in humans (Experiment 2), nor their behavior on the first benchmark (Experiment 1). The error scores are shown on the log scale.

We next used these measurements to create a benchmark that could be used to compare any computable object learning model – including the models considered in this study – against human object learning in this one-shot setting.

### 0.4 Models show weaker one-shot generalization compared to humans

As in Experiment 1, the benchmarking procedure for Experiment 2 consisted of 1) generating predictions of behavior from a model by having it perform the same experiment conducted in humans, then 2) scoring the similarity of that behavior to humans using an error statistic (MSE_*n*_). Thus, for each model, we replicated the one-shot behavioral experiment (n=16,000 simulated sessions per model), measured their accuracy on each of the 36 generalization tests described above, then compared those behavioral predictions to humans using MSE_*n*_.

Similar to our results from Experiment 1, here we found that models varied widely in their alignment with human learning behavior, and again found the top 1% subset of models achieved relatively low error (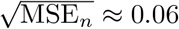, where the root noise floor is Approximately 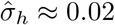). And as in Experiment 1, we found that *all* models had statistically significant differences in their behavior relative to humans, for this experiment. We also observed a positive relationship between the scores of the two benchmarks: models that were most human-like as evaluated by the benchmark based on Experiment 1 also tended to be the most human-like here, in the one-shot setting (see Fig 8D) – though *no* model explained human behavior in either experiment to the limits of statistical noise.

Part of the prediction failures we observed here lay in a failure to generalize as well as humans to several kinds of image variation. For example, we observed that all strong baseline models (identified from the benchmark from Experiment 1) had lower one-shot accuracy than humans in the presence of object pixel deletions, blur, shot noise, and scale shifts (see Fig 8C).

### 0.5 Specific individual humans outperform all models

Both benchmarks we developed in this study tested the ability of a model to predict human object learning at the “subject-averaged” level, where behavioral measurements drawn from several subjects are averaged together. This approach, by design, ignores any individual differences in learning behavior that may exist.

We wished to gauge the extent to which any such individual differences were present, and we performed an analysis on our behavioral data from Experiment 1. We identified subjects who performed all 64 subtasks in that experiment (22 out of 70 subjects total). We then attempted to reject the null hypothesis that there was no significant variation in their overall learning ability (see S3 Appendix). If this hypothesis were to be rejected, it would indicate that individuals must systematically vary in their learning behavior, at least in terms of their overall performance on these tasks. We indeed found that some subjects were reliably better object learners than others (*p* < 1e-4, permutation test).

Given this was the case, we next asked whether any of these individuals had an overall performance level higher than that of the highest performing model we identified in Experiment 1 (an encoding stage based on *ResNet152/avgpool*, and a tunable decision stage using the square plasticity rule). We identified n=5 individuals whose overall accuracy significantly exceeded that of this model (all p<1e − 5, Welch’s t-test, Bonferroni corrected). On average, this subset of humans had an overall accuracy of 0.92 *±* 0.01 (SEM over subjects); this was around 4% higher than this model’s average of 0.88.

## Discussion

A neurally mechanistic understanding of how humans accomplish visual object learning remains an open scientific problem. In this study, we focused on a core behavioral phenomenon entailed in visual object learning: the use of a finite number of image examples of new objects to accurately identify that object in new, unseen images. A necessary step in obtaining descriptions of the underlying neural mechanisms of this core phenomenon is evaluating the empirical alignment of alternative models with respect to measurements of human object learning behavior. To facilitate this, we first collected a set of human measurements in many tasks within this object learning setting (n=371,000 trials), allowing us to quantify the speed of human object learning (<10 trials to achieve close-to-asymptotic accuracy), the distinct pattern of learning difficulty they have for different objects, and their extent of generalization to specific image transformations after a single image example.

We then developed procedures to evaluate any image-computable object learning model over those same learning settings (which we refer to as “benchmarks” for human visual object learning), and tested a set of simple learning models (n=2,408 models) on those benchmarks. Each of these models consisted of two stages: 1) a fixed *encoding stage*, which maps incoming images to locations in an internal representational space, followed by 2) a tunable *decision stage* which aims to improve future choices by adjusting the plastic weights that convert that representation into a choice.

Prior to this study, we did not know if some or any of these learning models might be capable of explaining human object learning as assessed on naturalistic images like those used here. As such, we center our discussion on these models, but highlight that our raw behavioral data (and the associated behavioral benchmarks) are now a publicly available resource for testing image-computable object learning models beyond those evaluated here [GitHub].

### 0.6 Strengths and weaknesses of these object learning models

Linear learning on fixed image representations are strong baseline models of human object learning

On our first benchmark, which compares a learning model to humans under high view-variation learning conditions, we found a subset of models produced relatively accurate predictions of human learning behavior. The observed alignment of these models with humans does not originate from the fact they successfully learn new objects – these models also *fail* to rapidly learn the same objects that humans find difficult (Fig 6B), suggesting they have nontrivial similarities with humans, at least behaviorally.

We were surprised by the extent of similarity we observed between these models and humans, partly because some have suggested that DCNNs are unlikely to support adequate descriptions of human learning (e.g. [15, 39–42]), and partly because of the simplicity of these models. The results reported here suggest that learning models based on contemporary models of high-level visual neural representations and rudimentary, one-layer plasticity rules are a strong starting point to quantitatively account for the ability (and inability) of humans to learn arbitrary, new objects.

We note the present work does not directly engage the ongoing issue of whether the optimization mechanism of backpropagation is somehow involved in human learning [43]. Though the DCNN-based representations used in the learning models in this work were originally created using backpropagation, they were kept completely “frozen” over behavioral learning, with all behavioral learning achieved through a single layer of weight changes (which does not require backpropagation). At a conceptual level, this work regards DCNN-based representations as estimates of the (adult) human subject’s internal neural representation at the beginning of task learning, and is agnostic to how they were created before that point.

In general, the models we considered are composed only of operations that closely hew to those executed by first-order models of neurons – namely, linear summation of upstream population activity, ramping nonlinearities, and adjustment of local associational strengths using reward signals. This makes them not only plausible descriptions for the computations executed by the brain over object learning, but, with some additional assumptions (Fig 2B), they make predictions of neural phenomena.

For example, if the interpretation suggested in Fig 2B is taken at face value, these models make a couple of qualitative predictions. First, given the assumption that the encoding stage corresponds to the output of the ventral visual stream, these models predict that ventral stream representations used by humans over object learning need not undergo plastic changes to mediate behavioral improvements over the duration of the experiments we conducted (seconds-to-minutes timescale). This prediction is in line with prior studies showing adult ventral stream changes are typically moderate and take place on longer timescales (see [44] for review).

Moreover, the entirety of the computational learning mechanisms used by these models (i.e. the learning of thresholded, linear combinations of upstream neural activity using reward signals) can plausibly be executed at a single visuomotor synaptic interface where reward-based feedback signals are available. Several regions downstream of the ventral visual stream are possible candidates for this locus of plasticity during invariant object learning; we point to striatal regions receiving both high-level visual inputs and midbrain dopaminergic signals and involved in premotor processing, such as the caudate nucleus, as one set of candidates [45, 46].

#### Gaps between models and humans in few-shot learning

Despite the predictive strength of some models we tested, all models tested were unable to fully explain all replicable human behavior on either behavioral benchmark. One consistent prediction failure we observed in all models was a failure to learn new objects as rapidly as humans in low-sample regimes. We found this to be the case in both Experiment 1 (see Fig 7) and in Experiment 2, where we found that all tested models had lower accuracy than humans after one-shot across a variety of generalization tests (see Fig 8C). For example, we found that these models cannot one-shot generalize as well to scale shifts as humans, replicating previous work [47].

Taken together, these observations show all tested learning models currently have quantitative deficiencies from humans in the few-shot regime. We note that even if our experiments have underestimated human learning speed (e.g. from increased inattention rates on Mechanical Turk [48]), this inference would not change; the estimated gaps in few-shot learning abilities between these models and humans would be larger than the ones we report here. However, other aspects of similarity we found between models and humans – such as their shared patterns of relative difficulty – would be robust to such biases in our experiment.

### 0.7 Future visual object learning models to be tested

There are several potential ways to improve the predictive accuracy of the models we tested in this study (i.e. to find more human-like learning models). For example, it is possible that another model based on the conceptual model family we considered in this work could fully predict human learning over the benchmarks we developed, and we simply failed to implement and test that particular model here. If that is the case, such a model could differ from the ones we tested along one or both of its two components: its approximation of the visual representations used by humans during learning (i.e. its encoding stage), and/or its plasticity rule. Because the choice of plasticity rule had little effect on the predictive power of these models (Fig 5A) and did not interact significantly with the choice of encoding stage, we suggest it is more likely that alternative encoding stages would give rise to a more accurate model.

This view is consistent with the fact that the encoding stages we considered (Imagenet-pretrained DCNN representations) are known to only partially approximate the primate ventral stream, as directly measured by electrophysiological studies [36] and inferred by behavioral object categorization studies on images of already-learned objects [18, 32, 49]. If image-computable representations that more closely adhere to human visual representations could be built and/or identified, we anticipate they would lead to object learning models that close the prediction gap on the benchmarks we developed here.

Stepping back, it is also possible that *no* model from this conceptual model family could lead to fully accurate predictions on these benchmarks (or future benchmarks), but other types of models might do so. For example, one influential class of cognitive theories posits that the brain learns new objects by building structured, internal models of those objects from image exemplars, then uses those internal models to infer the latent content of each new image [8, 11, 13, 15, 16, 50]. It is possible that models based on these alternate approaches would generate more human-like learning over the tasks we tested here, and could be the key to achieving a full computational description of human object learning. In any case, implementing and testing these models on the benchmarks here is an important direction for future work.

### 0.8 Future extensions of object learning benchmarks

#### 0.8.1 Extensions of task paradigm

The two benchmarks we developed here certainly do not encompass all aspects of object learning. For example, each benchmark focused on discrimination learning between two novel objects, but humans can potentially learn and report on many more objects simultaneously. Moreover, humans can readily learn object *categories* at different levels of abstraction, each of which may encompass multiple specific objects [51]. The models tested here scale naturally to task paradigms involving additional objects (via the incorporation of new linear choice preferences to the decision stage), and are capable of learning categories of varying abstraction; comparing them to humans in those richer learning settings (and identifying any of their limits in those settings) could strongly motivate the consideration of more complex models.

#### 0.8.2 Extending stimulus presentation time

For presenting stimuli, we followed conventions used in previous visual neuroscience studies [18, 52] of object perception: achromatic images containing single objects rendered with high view uncertainty on random backgrounds, presented at <10 degrees of visual field and for <200 milliseconds.

The chosen stimulus presentation time of 200 milliseconds is too short for a subject to initiate a saccadic eye movement based on the content of the image [53]. Such a choice simplifies the input of any model (i.e., to a single image, rather than the series of images induced by saccades); on the other hand, active viewing of an image via target-directed saccades might be a central mechanism deployed by humans to mediate learning of new objects.

We note that if this is the case, our task paradigm (which would disrupt any such saccade-based mechanisms from being used) would be underestimating the number of images needed by humans to achieve learning on new objects, compared to a scenario in which subjects had unlimited viewing time on each trial. Thus, removing such a bias in our experimental design could potentially reveal larger differences between models and humans.

Beyond extending viewing time, designing tasks which more closely hew to typical object learning contexts for humans (e.g. involving colored images and/or movies of potentially multiple objects physically embedded in natural scenes) will be an important direction for future work.

#### 0.8.3 Differences between individual subjects

We primarily focused on studying human learning at the subject-averaged level, where behavioral measurements are averaged across several individuals (i.e. subject-averaged learning curves; see Fig 1D). However, individual humans may have systematic differences in their learning behavior that are (by design) ignored with this approach.

For example, we found that individual subjects may differ in their overall learning abilities: we identified a subpopulation of humans who were significantly more proficient at learning compared to other humans (see Fig 9B). We did not attempt to model this individual variability in this study; whether these differences can be explained by alterations to this model family, if at all, (e.g. through the introduction of random effects to the parameters of the encoding stage and/or plasticity rules) remains an area for future study.

**Fig 9.**
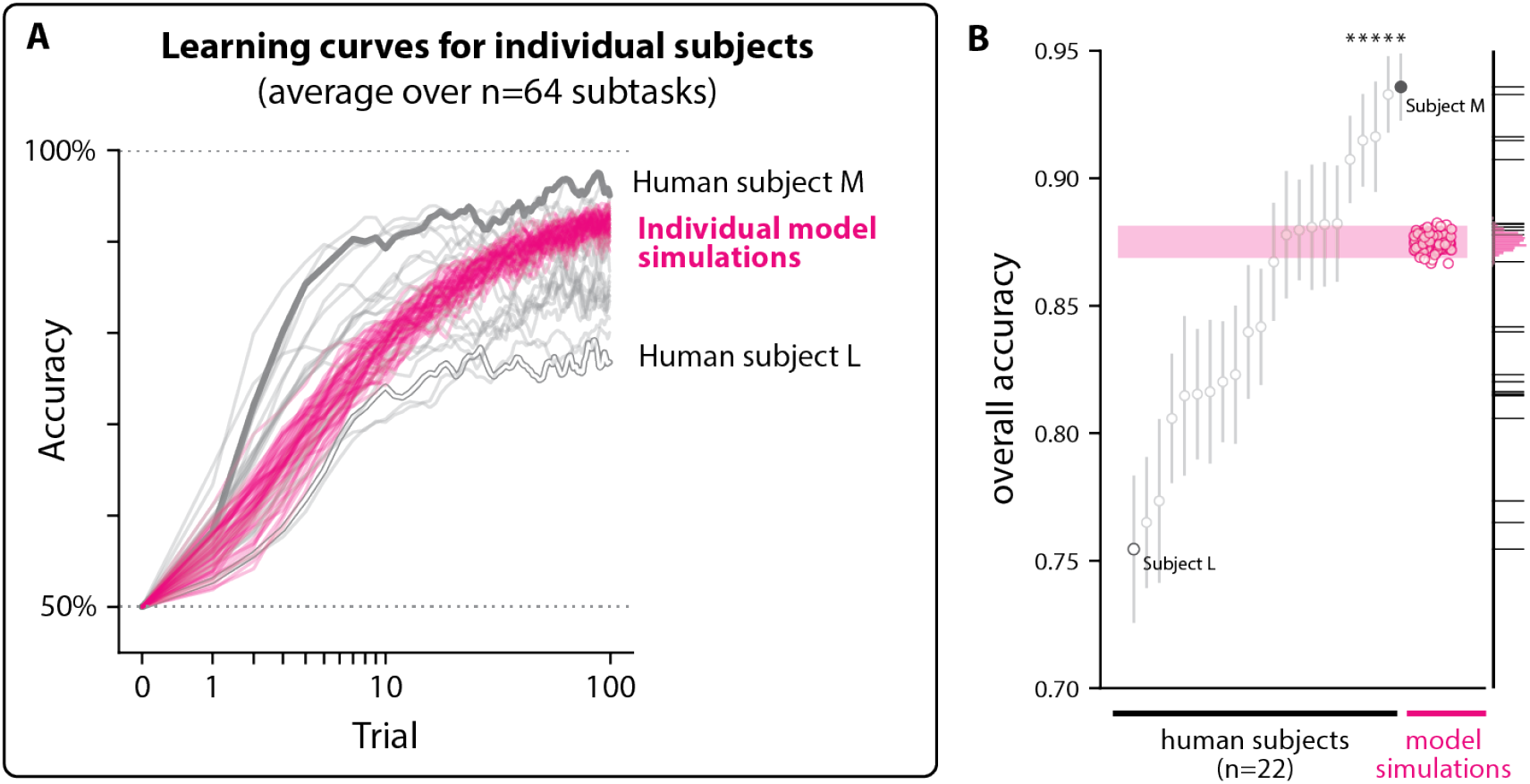
Individual differences in learning ability. **A. Individual-level learning curves**. We identified 22 subjects who performed all 64 subtasks in Experiment 1, and computed their subtask-averaged learning curves. Each gray curve corresponds to the learning curve for a different individual subject (smoothed using state-space estimation from [38]). In humans, a range of overall learning performance is seen: some subjects consistently outperformed others (e.g. Subject M, highest accuracy over all trials and subtasks), while others consistently underperformed (e.g. Subject L, lowest average accuracy). In magenta are subtask-averaged learning curves corresponding to individual model simulations from the highest-performing model e tested in this study (encoding stage = ResNet152/avgpool, plasticity rule = square). **B. Some individual humans outperform all models**. Five out of 22 subjects had significantly higher overall performance than the highest performing model we tested (one-tailed Welch’s t-test, Bonferroni corrected, p<0.05).

Moreover, performing subject-averaging is known to lead to the masking of learning dynamics present at the level of single subjects, such as delayed rises or step increases in accuracy [54]. Performing analyses to compare any such learning dynamics between individual humans and learning models is another important extension of our work.

Lastly, we did not attempt to model any systematic increases in a subject’s learning performance as they performed more and more subtasks available to them (in either Experiment 1 or 2). This phenomenon (learning-to-learn, learning sets, or meta-learning) is well-known in psychology [55], but to our knowledge has not been systematically measured or modeled in the domain of human object learning. Expanding these benchmarks (and models) to measure and account for such effects is an important future step of building models of the work done here.

## Materials and methods

### 0.9 Overview of experiments

For both experiments, the core measurement we sought to obtain was the discrimination performance of a typical subject as they received increasing numbers of exposures to images of the to-be-learned (i.e. new) objects.

We assumed that different pairs of objects result in potentially different rates of learning, and we wanted to capture those differences. Thus, in Experiment 1, we aimed to survey the empirical landscape of this human ability by acquiring this learning curve measurement for many different pairs of objects (n=64 pairs). Specifically, for each pair of to-be-learned objects (referred to as a “subtask”), we aimed to measure (subject-averaged) human learning performance across 100 learning trials, where each trial presented a test image generated by one of the objects under high viewpoint uncertainty (e.g. random backgrounds, object location, and scale). We refer to this 100-dimensional set of measurements as the *learning curve* for each subtask.

In Experiment 2, we aimed to measure the pattern of human learning that results from their experience with just a *single* canonical example of each of the to-be-learned objects (a.k.a. “one-shot learning”). Specifically, we wished to measure the pattern of human discrimination ability over various kinds of identity-preserving image transformations (e.g, object scaling, transformation, and rotation). In total, we tested nine kinds of transformations. We anticipated that humans would show distinct patterns of generalization across these transformations, and we aimed to measure the human commonalities in those patterns (i.e. averages across subjects).

Experiments 1 and 2 both utilized a two-way object learning task paradigm that is conceptually outlined in Fig 1B. The two experiments differed only in the manner in which test images were generated and sampled for presentation, and we describe those differences in detail in their respective sections. Before that, we provide more detail on the specific procedures and parameters we used to implement the common two-way object learning task paradigm.

### 0.10 Task paradigm

For both experiments, human subjects were recruited from Mechanical Turk [56], and ran tasks on their personal computers. Demographic information (age, sex, gender, or ethnicity) was not collected; all online subjects were anonymous. We also recruited *n* = 4 subjects for testing in the lab. We designed and administered these experiments in accordance to a protocol approved by the Massachusetts Institute of Technology Committee on the Use of Humans as Experimental Subjects (Protocol # 0812003043A017).

Each experiment (Experiments 1 & 2) consisted of a set of subtasks. For each subtask, we asked a population of human subjects to learn that subtask, and we refer to the collection of trials corresponding to a specific subject in a subtask as a “session”.

At the beginning of each session, the subject was instructed that there would be two possible objects – one belonging to the “F” category and the other belonging to the “J” category. The subject’s goal was to correctly indicate the category assignment for each test image. The specific instructions were: *“On each trial, you’ll view a rapidly flashed image of an object. Your task is to figure out which button to press (either “F” or “J” on your keyboard) after viewing a particular image. Each button corresponds to an object (for example, a car might correspond to F, while a dog might correspond to J)*.*”*

Subjects were also informed that they would receive a monetary bonus (in addition to a base payment) for each correctly indicated test image, incentivizing them to learn. We next describe the structure of a single trial in detail below.

#### 0.10.1 Test image presentation

Each trial began with a display start screen that was uniformly gray except for a small black dot at the center of the screen, which indicated the future center of each test image.^2^ We intended for this fixation point to encourage the subject to consistently view each test image at the center of their field of view (see S2 for in-lab eye measurements). The subject then initiated the trial by pressing the space bar on their keyboard. Once pressed, a test image (occupying ≈6*°* of the visual field) belonging to one of the two possible object categories immediately appeared. That test image remained on the screen for ≈ 200 milliseconds before disappearing (and returning the screen to uniform gray).^3^

For each subject and each trial, the test image was selected by first randomly picking (with equal probability) one of the two objects as the generator of the test image. Then, given that selected object, an image of that object was randomly selected from a pool of pre-rendered possible images. Test images were always selected without replacement (i.e. once selected, that test image was removed from the pool of possible future test images for that behavioral session).

#### 0.10.2 Subject choice reporting

Fifty milliseconds after the disappearance of the test image, the display cued the subject to report the object that was “in” the image. The display showed two identical white circles – one on the lower left side of the fixation point and the other on the lower right side of the fixation point. The subject was previously instructed to select either the “F” or “J” keys on their keyboard. We randomly selected one of the two possible object-to-key mappings prior to the start of each session, and held it fixed throughout the entire session. This mapping was not told to the subject; thus, on the first trial, subjects were (by design) at chance accuracy.

To achieve perfect performance, a subject would need to associate each test image of an object to its corresponding action choice, and not to the other choice (i.e., achieving a true positive rate of 1 and a false positive rate of 0).

Subjects had up to 10 seconds to make their choice. If they failed to make a selection within that time, the task returned to the trial initiation phase (above) and the outcome of the trial was regarded as being equivalent to the selection of the incorrect choice.^4^.

#### 0.10.3 Trial feedback

As subjects received feedback which informed them whether their choice was correct or incorrect (i.e. corresponding to the object that was present in the preceding image or not), they could in principle learn object-to-action associations that enabled them to make correct choices on future trials.

Trial feedback was provided immediately after the subject’s choice was made. If they made the correct choice, the display changed to a feedback screen that displayed a reward cue (a green checkmark). If they made an error, a black “x” was displayed instead. Reward cues remained on the screen for 50 milliseconds, and were accompanied by an increment to their monetary reward (see above). Error cues remained on the screen for 500 milliseconds. Following either feedback screen, a 50 millisecond delay occurred, consisting of a uniform gray background. Finally, the display returned to the start screen, and the subject was free to initiate the next trial.

### 0.11 Experiment 1: Learning objects under high view variation

Our primary human learning benchmark (Experiment 1) was based on measurements of human learning curves over subtasks involving images of novel objects rendered under high view-variation. We describe our procedure for generating those images, collecting human behavioral measurements, and benchmarking models against those measurements below.

#### 0.11.1 High-variation stimulus image generation

We designed 3D object models (n=128) using the “Mutator” generative design process [57]. We generated a collection of images for each of those 3D objects using the POV-Ray rendering program [58]. To generate each image, we randomly selected the viewing parameters of the object, including its projected size on the image plane (25%-50% of total image size, uniformly sampled), its location (*±*40% translation from image center for both *x* and *y* planes, uniformly sampled), and its pose relative to the camera (uniformly sampled random 3D rotations). We then superimposed this view on top of a random, naturalistic background drawn from a database used in a previously reported study [52]. All images used in this experiment were grayscale, and generated at a resolution of 256×256 pixels. We show an example of 32 objects (out of 128 total) in Fig 1A, along with example stimulus images for two of those objects on the right.

#### 0.11.2 Design of subtasks

We randomly paired the 128 novel objects described above into pairs (without replacement) to create n=64 subtasks for Experiment 1, each consisting of a distinct pair of novel objects. Each behavioral session for a subtask consisted of 100 trials, regardless of the subject’s performance. On each trial of a session, one of the two objects was randomly selected, and then a test image of that object was drawn randomly without replacement from a pre-rendered set of 100 images of that object (generated using the process above). That test image was then presented to the subject (as described in Test image presentation). We collected 50 sessions per subtask and all sessions for each subtask were obtained from separate human subjects, each of whom we believe had not seen images of either of the subtask’s objects before participation.

#### 0.11.3 Subject recruitment and data collection

Human subjects were recruited on the Mechanical Turk platform [56] through a two-step screening process. The goal of the first step was to verify that our task software successfully ran on their personal computer, and to ensure our subject population understood the instructions. To do this, subjects were asked to perform a prescreening subtask with two common objects (elephant vs. bear) using 100 trials of the behavioral task paradigm (described in Task paradigm above). If the subject failed to complete this task with an average overall accuracy of at least 85%, we intentionally excluded them from all subsequent experiments in this study.

The goal of the second step was to allow subjects to further familiarize themselves with the task paradigm. To do this, we asked subjects to complete a series of four “warmup” subtasks, each involving two novel objects (generated using the same “Mutator” software, but distinct from the 128 described above). Subjects who completed all four of these warmup subtasks, regardless of accuracy, were enrolled in Experiment 1. Data for these warmup subtasks were not included in any analysis presented in this study. In total, we recruited n=70 individual Mechanical Turk workers for Experiment 1.

Once a subject was recruited (above), they were allowed to perform as many of the 64 subtasks as they wanted, though they were not allowed to perform the same subtask more than once (median n=61 total subtasks completed, min=1, max=64). We aimed to measure 50 sessions per subtask (i.e. 50 unique subjects), where each subject’s session consisted of an independently sampled, random sequence of trials. Each of these subtasks followed the same task paradigm (described in Methods Task paradigm), and each session lasted 100 trials. Thus, the total amount of data we aimed to collect was 64 subtasks *×* 100 trials *×* 50 subjects = 320*k* measurements.

#### 0.11.4 Behavioral statistics in humans

We aimed to estimate a typical subject’s accuracy at each trial, conditioned on a specific subtask. We therefore computed 64 *×* 100 accuracy estimates (*subtask × trial*) by taking the sample mean across subjects. We refer to this [64, 100] matrix of point statistics as *Ĥ*. Each row vector *Ĥ*_*s*_ has 100 entries, and corresponds to the mean human “learning curve” for subtask *s* = {1, 2, …64}.

Because each object was equally likely to be shown on any given test trial, each of these 100 values of *Ĥ*_*s*_ may be interpreted as an estimate of the average of the true positive and true negative rates (i.e. the balanced accuracy). The balanced accuracy is related to the concept of *sensitivity* from signal detection theory – the ability for a subject to discriminate two categories of signals [59]. We note that an independent feature of signal detection behavior is the *bias* – the prior probability with which the subject would report a category. We did not attempt to quantify or compare the bias in models and humans in this study.

#### 0.11.5 Simulating behavioral sessions in computational models

To obtain the learning curve predictions of each computational model, we required that each model perform the same set of subtasks that the humans performed, as described above. We imposed the same requirements on the model as we did on the human subjects: that it begins each session without knowledge of the correct object-action contingency, that it should generate a action choice based solely on a pixel image input, and that it can update its future choices based on the history of scalar-valued feedback (“correct” or “incorrect”). If the choices later in the session are more accurate than those earlier in the session, then we colloquially say that the model has “learned”, and comparing and contrasting the learning curves of models with those of humans was a key goal of Experiment 1.

We ran n=32,000 simulated behavioral sessions for each model (500 simulated sessions for each of the 64 subtasks), where on each simulation a random sequence of trials was sampled in an identical fashion as in humans (see above). During each simulation, we recorded the same raw “behavioral” data as in humans (i.e. sequences of correct and incorrect choices), then applied the same procedure we used to compute *Ĥ* (see above) to compute an analogous collection of point statistics on the model’s raw behavior, which we refer to as 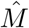.

#### 0.11.6 Comparing model learning with human learning

The learning behavior generated by an computable model of human learning 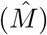 should minimally replicate the measured learning behavior of humans (i.e.*Ĥ*), to the limits of statistical noise. To identify any such models, we developed a scoring procedure to compare the similarity of the learning behavior in humans with any candidate learning model. We describe this procedure below.

##### Bias-corrected mean squared error

Given a collection of human measurements *Ĥ* (here, a matrix of accuracy estimates for *S* = 64 subtasks over *T* = 100 trials) and corresponding model measurements 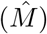, we computed a standard goodness-of-fit metric, the mean-squared error (MSE; lower is better). The formula for the MSE is given by:

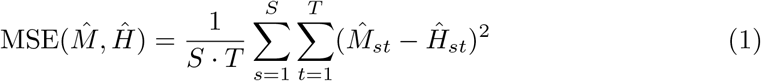

Because *Ĥ*_*st*_ and 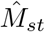 are random variables (i.e. sample means), the MSE itself is a random variable. It can be seen that the expected value of 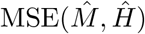 consists of two conceptual components: the expected difference between the model and humans, and *noise components*:

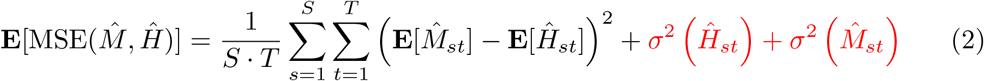

Where **E**[.] denotes the expected value, and *σ*^2^(.) denotes the variance due to finite sampling (a.k.a. “noise”).

Equation (2) shows that the expected MSE for a model depends not only on its expected predictions 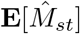, but also its sampling variance 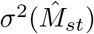. In the present case where 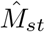 is the mean over independent (but not necessarily identically distributed) Bernoulli variables, the value of 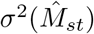 happens to depend on the expected prediction of the model itself, 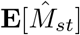. ^5^

Because the sampling variance of the model depends on its predictions, it is therefore conceptually possible that a model with worse (expected) predictions could achieve a lower expected MSE, simply because its associated sampling variance is lower.^6^

We corrected for this inferential bias by estimating, then subtracting, these variance terms from the “raw” MSE for each model we tested.^7^. We refer to this bias-corrected error as MSE_*n*_.

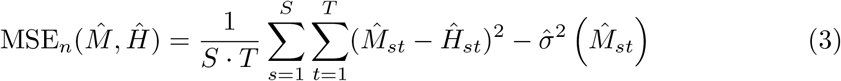

Where 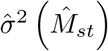 is an unbiased estimator of the variance of 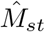. We write the equation for 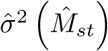 below, where *k*_*st*_ is the number of observed correct choices over the *n*_*st*_ model simulations conducted for subtask *s* and trial *t*:

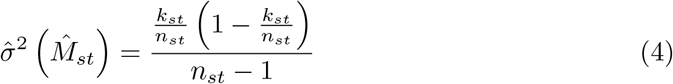

Because 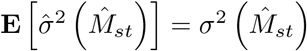, the expected value of MSE_*n*_ can be shown to be:

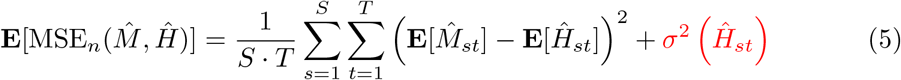

Intuitively, MSE_*n*_ is an estimate of the mean-squared error that would be achieved by a model if we had a noiseless estimate of its predictions (i.e. had an infinite number of simulations of that model been performed). We note that the value of its square root, 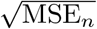, gives a rough^8^ estimate of the average deviation between a model’s prediction and human measurements, in units of the measurements (in this study, units of accuracy).

##### Noise floor estimation

It can be in Equation (5) that there are terms *σ*^2^(*Ĥ*_*st*_), originating from the uncertainty in our experimental estimates of human behavior. These terms are always positive, and create a lower bound on the expected MSE_*n*_ for all models. That is, even if a model is expected to perfectly match the subject-averaged behavior of humans (where **E**[*M*_*st*_] = **E**[*H*_*st*_], for all subtasks *s* and trials *t*), it cannot be expected to achieve an error below this lower bound. We call this lower bound the “noise floor”, and use the symbol 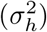 to refer to it:

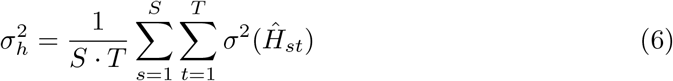

It is possible to make an unbiased estimate of the noise floor 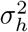 if one can make unbiased estimates of each 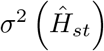 term. We did so by using the unbiased estimator from Equation (4). We write the full expression for our estimate of the noise floor, 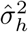, below:

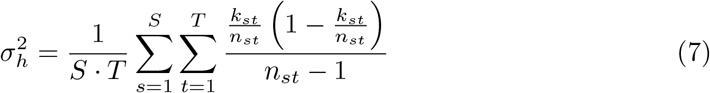

Where *k*_*st*_ is the number of human subjects (out of *n*_*st*_ total subjects) that made a correct choice on subtask *s* and trial *t*. The square root of this value, *σ*_*h*_, gives a rough estimate of the average deviation one would expect in our subject-averaged measurements of behavior (*Ĥ*_*st*_) over repetitions of the experiment (i.e. upon another resampling of human subjects and behavioral sessions).

##### Null hypothesis testing

For each model we tested, we attempted to reject the null hypothesis that 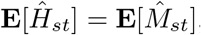, for all subtasks *s* and trials *t*. To do so, we first approximated the distribution for 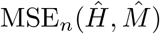 that would be expected under this null hypothesis, using bootstrapping.

To do so, we first computed bootstrap replicates of *Ĥ* and approximated samples of the null model 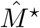 (where 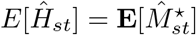). A bootstrap replicate of *Ĥ* was constructed by first resampling individual human sessions without replacement, taking the same number of resamples per subtask as in the original experiment. We then computed the replicate 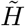 using the same procedure described in Behavioral statistics in humans. Behavior from the null model cannot be sampled directly (i.e. we do not have the “true model” of human learning), but by definition shares the same expected behavior as a randomly sampled, individual human. We therefore created a bootstrap sample of the null model 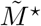 by (also) taking resamples of individual human sessions, setting the number of resamples per subtask to the number of model simulations conducted per subtask (here, n=500 simulations per subtask). We then computed and saved 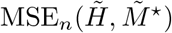 for that iteration, and repeated this process for B=1,000 iterations to obtain an approximate null distribution for MSE_*n*_.

If a model’s actual 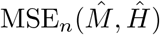 score fell above the *α*-quantile of the estimated null distribution, we rejected it on the basis of having significantly more error than what would be expected from a “true” model of humans (with estimated significance level *α*).

##### Lapse rate correction

Lastly, we corrected for any lapse rates present in the human data. We defined the lapse rate as the probability with which a subject would randomly guess on a trial, and we assumed this rate was constant across all trials and subtasks. To correct for any such lapse rate in the human data, we fit a simulated lapse rate *γ* parameter to each model, prior to computing its MSE_*n*_. Given a lapse rate parameter of *γ* (ranging between 0 and 1), a model would, on each trial, guess randomly with probability *γ*. For each model, we identified the value of *γ* that minimized its empirical MSE_*n*_.

We note that fitting *γ* can only drive the behavior of a model toward randomness; it cannot artificially introduce improvements in its learning performance.

### 0.12 Experiment 2: One-shot human object learning benchmark

For the second benchmark in this study, we compared one-shot generalization in humans and models. Our basic approach was to allow humans to learn to distinguish between two novel objects using a single image per object, then test them on new, transformed views of the support set.

#### 0.12.1 One-shot behavioral testing

We used the same task paradigm described in Task paradigm (i.e. two-way object discrimination with evaluative feedback). We created 64 object models for this experiment (randomly paired without replacement to give a total of 32 subtasks). These objects were different from the ones used in the previous benchmark (described in Experiment 1: Learning objects under high view variation).

At the beginning of each session, we randomly assigned the subject to perform one of 32 subtasks. Identical to Experiment 1, each trial required that the subject view an image of an object, make a choice (“F” or “J”), and receive feedback based on their choice. Each session consisted of 20 trials total, which was split into a “training phase” and “testing phase”, which we describe below.

##### Training phase

The first ten trials (the “training phase”) of the session were based on a single image for each object object (i.e. n=2 distinct images were shown over the first 10 trials). We ensured the subject performed trial with each training image five times total in the training phase; randomly permuting the order in which these trials were shown.

##### Testing phase

On trials 11-20 of the session (the “testing phase”), we presented trials containing new, transformed views of the two images used in the training phase. For each trial in the test phase, we randomly sampled an unseen test image, each of which was a transformed version of one of the training images. There were 36 possible transformations (9 transformation types, with 4 possible levels of strength). We describe how we generated each set of test images in the next section (see Fig 1B for examples). On the 15th and 20th trial, we presented “catch trials” consisting of the original training images. Throughout the test phase, we continued to deliver evaluative feedback on each trial.

#### 0.12.2 One-shot stimulus image generation

Here, we describe how we generated all of the images used in Experiment 2. First, we generated each 3D object model using the Mutator process (see High-variation stimulus image generation). Then, for each object (n=64 objects), we generated a single canonical training image – a 256×256 grayscale image of the object occupying ≈50% of the image plane, centered on a gray background. We randomly sampled its three axes of pose from the uniform rotational distribution.

For each training image, we generated a corresponding set of test images by applying different kinds of image transformations we wished to measure human generalization on.

In total, we generated test images based on 9 transformation types, and we applied each transformation type at 4 levels of “strength”. We describe those 9 types with respect to a single training image, below.

##### Translation

We translated the object in the image plane of the training image. To do so, we randomly sampled a translation vector in the image plane (uniformly sampling an angle from *θ* ∈ [0°, 360°]), and translated it *r* pixels in that direction. We repeated this process (independently sampling *θ* each time) for *r* = 16, 32, 64, and 96 pixels (where the total image size 256 × 256 pixels), for two iterations (for a total of eight translated images).

##### Backgrounds

We gradually replaced the original, uniform gray background with a randomly selected, naturalistic background. Each original background pixel *b*_*ij*_ in the training image was gradually replaced with a naturalistic image *c* using the formula 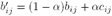. We varied *α* at four logarithmically spaced intervals, *α* = 0.1, 0.21, 0.46, 1. Note that at *α* = 1, the original gray background is completely replaced by the new, naturalistic background. We generated two test images per *α* level, independently sampling the background on each iteration (for a total of eight images per object).

##### Scale

We rescaled the object’s size on the image to 12.5%, 25%, 50%, and 150% of the original size (four images of the object at different scales).

##### Out-of-plane rotations

We rotated the object along equally spaced 45° increments, rendering a test image at each increment. We did so along two separate rotational axes (horizontal and vertical), leading to n=13 test images total based on out-of-plane rotations.

##### In-plane rotation

We rotated the object inside of the image-plane, along 45° increments. This resulted in n=7 test images based on in-plane rotations.

##### Contrast

We varied the contrast of the image. For each pixel *p*_*ij*_ (where pixels range in value of 0 and 1), we adjusted the contrast using the equation 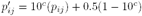, varying *c* from −0.8, −0.4, 0.4 and 0.8.

##### Pixel deletion

We removed pixels corresponding to the object in the training image, replacing them with the background color (gray). We removed 25%, 50%, 75%, and 95% of the pixels, selecting the pixels randomly for each training image.

##### Blur

We blurred the training image using a Gaussian kernel. We applied blurring with kernel radii of 2, 4, 8, and 16 pixels (with an original image resolution of 256 *×* 256 pixels) to create a total of 4 blurred images.

##### Gaussian noise

We applied Gaussian noise to the pixels of the training image. For each pixel *p*_*ij*_, we added *i*.*i*.*d*. Gaussian noise:

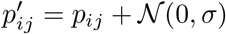

We applied noise with *σ* = 0.125, 0.25, 0.375 and 0.5 (where pixels range in luminance value between 0 and 1). We then clipped the resultant pixels to lie between 0 and 1.

#### 0.12.3 Human behavioral measurements for Experiment 2

Subject recruitment

We used the same two-step subject recruitment procedure described above (see Subject recruitment and data collection), and recruited n=170 human subjects. Some of these subjects overlapped with those in Experiment 1 (n=9 subjects participated in both experiments).

All recruited subjects were invited to participate in up to 32 behavioral sessions. We disallowed them from repeating subtasks they had performed previously. Subjects were required to perform a minimum of four such behavioral sessions. In total, we collected n=2,547 sessions (≈ 51k trials) for Experiment 2.

##### One-shot behavioral statistics in humans

We aimed to estimate the expected accuracy of a subject on each of the 36 possible transformations, correcting for attentional and memory lapses.

To do so, we combined observations across the eight test trials in the testing phase to compute accuracy estimate for each of the 36 transformations; that is, we did not attempt to quantify how accuracy varied across the testing phase (unlike the previous benchmark). We also combined observations across the 32 subtasks in this experiment. In doing so, we were attempting to measure the *average* generalization ability for each type of transformation (at a specific magnitude of transformation change from the training image), ignoring the fact that generalization performance likely depends on both the objects to be discriminated (i.e. the appearance of the objects in each subtask), the specific training images that were used, and the testing views of each object (e.g. the specific way in which an object was rotated likely affects generalization – not just the absolute magnitude of rotation). In total, we computed 36 point statistics (one per transformation).

##### Estimating performance relative to catch performance

Here we assumed that each human test performance measurement was based on a combination of the subject’s ability to successfully generalize, a uniform guessing rate (i.e. the probability with which a subject executes a 50-50 random choice), and the extent to which the subject successfully acquired and recalled the training image-response contingency (i.e. from the first 10 trials). We attempted to estimate the test performance of a human subject that could 1) fully recall the association between each training image and its correct choice during the training phase, and 2) had a guess rate of zero on the test trials.

To do so, we used trials 15 and 20 of each session, where one of the two training images was presented to the subject (“catch trials”). Our main assumption here was that performance on these trials would be 100% assuming the subject had perfect recall, and had a guess rate of zero. Under that assumption, the actual, empirically observed accuracy *p*_catch_ would be related to any overall guess and/or recall failure rate *γ* by the equation *γ* = 2 − 2*p*_catch_. We then adjusted each of the point statistics (i.e. test performances) to estimate their values had *γ* been equal to zero, by applying the following formula:

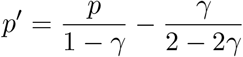

We refer to the collection of 36 point statistics (following lapse rate correction) as *Ĥ*^*os*^.

#### 0.12.4 Comparing model one-shot learning with human one-shot learning

Model simulation of Experiment 2

For this benchmark, we required that a model perform a total of 16,000 simulated behavioral sessions (500 simulated sessions for each of the 32 possible subtasks). Each simulated session proceeded using the same task paradigm as in humans (i.e. 10 training trials, followed by a test phase containing 8 test trials and 2 catch trials). Based on the model’s behavior over those simulations, we computed the same set of point statistics described above, though we did not correct for any attentional lapses or recall lapses in the model, which we assumed was absent in models. In this manner, for each model, we obtained a collection of point statistics reflecting their behavior on this experiment, 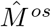.

#### Bias-corrected mean-squared error and null hypothesis testing

We used the same statistical approach for our primary benchmark (introduced in Comparing model learning with human learning) to summarize the alignment of a model with humans. That is, we used the bias-corrected error metric MSE_*n*_ as our metric of comparison:

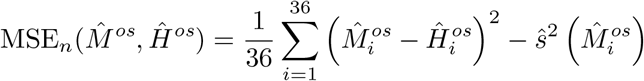

We estimated the null distribution for MSE_*n*_ using bootstrap resampling, following the same procedure outlined in the first benchmark (bootstrap-resampling individual sessions).

### 0.13 Baseline model family

For a model to be scored on the benchmarks we described above, it must fulfill only the following three requirements: 1) it takes in any pixel image as its only sensory input (i.e. it is image computable), 2) it can produce an action in response to that image, and 3) it can receive scalar-valued feedback (rewards). Here, we implemented several models which fulfill those requirements.

All models we implemented consist of two components. First, there is an *encoding stage* which re-represents the raw pixel input as a vector 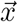 in a multidimensional Euclidean space.The parameters of this part of the model are held fixed (i.e., no learning takes place in the encoding stage).

The second part is a *tunable decision stage*, which takes that representational vector and produces a set of *C* choice preferences (in this study, *C* = 2). The preference for each choice is computed through a dot product 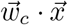, where 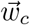 is a vector of weights for choice *c*. The choice with the highest preference score is selected, and ties are broken randomly.

After the model makes its choice, the environment may respond with some feedback (e.g. positive or negative reward). At that point, the decision stage can process that feedback and use it to change its parameters (i.e. to learn). All learning in the models tested here takes place only in the parameters of the decision stage (all weight vectors 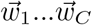); the encoding stage has completely fixed parameters.

In total, any given model in this study is defined by these two components – the encoding stage and the decision stage. We provide further details for those two components below.

#### 0.13.1 Encoding stages

The encoding stages were intermediate layers of deep convolutional neural network architectures (DCNNs). We drew a selection of such layers from a pool of 19 network architectures available through the PyTorch library [60], each of which had pretrained parameters for solving the Imagenet object classification task [37].

For each architecture, we selected a subset of these intermediate layers to test in this study, spanning the range from early on in the architecture to the final output layer (originally designed for Imagenet). We resized pixel images to a standard size of 224×224 pixels using bilinear interpolation. In total, we tested n=344 intermediate layers as encoding stages.

##### Dimensionality reduction

Once an input image is fed into a DCNN architecture, each of its layers produces a representational vector of a dimensionality specified by the architecture of the model. Depending on the layer, this dimensionality may be relatively large (¿10^5^), making it hard to efficiently perform numerical calculations on contemporary hardware. We therefore performed dimensionality reduction as a preprocessing step. We performed dimensionality reduction using random Gaussian projections to a standard size of 2048, if the original dimensionality of the layer was greater than this number. This procedure approximately preserves the original representational structure of the layer (i.e., pairwise distances between points in that space) [61] and is similar to to computing and retaining the first 2048 principal components of the representation.

##### Feature normalization

Once dimensionality reduction was performed, we performed another standardization step. We computed centering and scaling parameters for each layer, so that its activations fit inside a sphere of radius 1 centered about the origin (i.e. 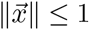, for all 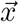).

To do so, we computed the activations of the layer over using the images from the “warmup” tasks human subjects were exposed to prior to performing any task in this study (i.e. 50 randomly selected images of 8 objects, see Subject recruitment and data collection). We computed the sample mean of those activations, and set this as the new origin of the encoding stage (i.e. the centering parameter). Then, we took the 99th quantile of the activation norms (over those same images) to calculate the approximate radius of the representation, and set this as our scaling parameter (i.e. dividing all activations by this number). Any activations with a norm greater than this radius were scaled to have a norm of 1.

Other kinds of feature standardization schemes are possible: for instance, one could center and scale the sensory representations for each subtask separately. However, such a procedure would expose models to the statistics of subtasks that are meant to be independent tests of their ability to learn new objects – statistics which we considered to be predictions of the encoding stage.

#### 0.13.2 Tunable decision stage

Once the encoding stage re-represents an incoming pixel image as a multidimensional vector 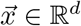, a *tunable decision stage* takes that vector as an input, and produces a choice as an output.

##### Generating a decision

To select a choice, the tunable decision stage first generates choice preferences for each of the *C* possible actions, using the dot products 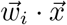 for *i* = 1…*C*. Then, the choice with the highest preference is selected (*c* = **argmax**_*i*_ 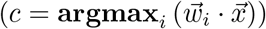). If all choices have the same preference value, a choice is randomly selected.

##### Learning from feedback

Once a choice is selected, the environment may convey some scalar-valued feedback (e.g. reward or punish signals). The model may use this feedback to change its future behavior (i.e., to learn). For all models considered here, this may be accomplished (only) by changing its weight vectors 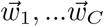. Thus, learning on each trial can be summarized by the changes to each weight vector by some 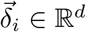:

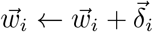

There are many possible choices on how each 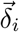 may be computed from feedback; here, we focused on a set of seven rules based on the stochastic gradient descent algorithm for training a binary classifier or regression function. In all cases except one,^9^ the underlying strategy of each decision stage can be understood as *predicting the reward* following the selection of each possible choice, and using these predictions to select the choice it believes will lead to the highest reward.

Specifically, we tested the plasticity rules induced by the gradient descent update on the perceptron, cross-entropy, exponential, square, hinge, and mean absolute error loss functions (shown in Fig 2D), as well as the REINFORCE plasticity rule. They are summarized in Table 0.13.2.

Each of these plasticity rules has a single free parameter – a learning rate. For each plasticity rule, there is a predefined range of learning rates that guarantees the non-divergence of the decision stage, based on the smoothness or Lipschitz constant of each of the plasticity rule’s associated loss function [62]. We did not investigate different learning rates in this study; instead, we simply selected the highest learning rate possible (such that divergence would not occur) for each plasticity rule.

## Supporting information

### S1 Table. Summary of plasticity rules

Each plasticity rule can be understood by the update 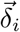 it generates for each weight vector 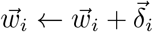, based on the current input 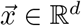, the selected choice *c* ∈ {0, 1, …*C*}, and the subsequent environmental reward *r* ∈ [−1, 1]. Each plasticity rule is parameterized by a learning rate *α*, between 0 and 1. These equations assume that the input 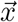 has a bounded norm of 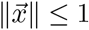.

### S1 Appendix. Effect of model choices on human behavioral similarity

As described in Section 0.13, each model in this study was defined by two components (the encoding stage and the plasticity rule). We wished to evaluate the effect of each of these components in driving the similarity of the model to human behavior. For example, it was possible that all models with the same encoding stage had the same learning score, regardless of which plasticity rule they used (or vice versa).

To test for these possibilities, we performed a two-way ANOVA over all observed model scores (in MSE_*n*_) computed in this study, using the encoding stage and plasticity rule as the two factors, and MSE_*n*_ as the dependent variable. By doing so, we were able to estimate the amount of variation in model scores that could be explained by each individual component, and thereby gauge their relative importance. We briefly describe the procedure for this analysis below. First, we wrote the MSE_*n*_ score of each model as a combination of four variables:

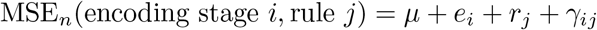

Where *μ* is the average MSE_*n*_ score, over all models. The variables *e*_*i*_ and *r*_*j*_ encode the value of the average difference from *μ* given encoding stage *i* and rule *j*, respectively.

| Plasticity rule | Weight update for selected choice ($\vec{\delta}_c$) |
| --- | --- |
| Perceptron | $\vec{\delta}_c = \begin{cases} \alpha r \vec{x} & \text{if } r(\vec{w}_c \cdot \vec{x}) \leq 0 \\ 0 & \text{else} \end{cases}$ |
| Hinge | $\vec{\delta}_c = \begin{cases} \alpha r \vec{x} & \text{if } r(\vec{w}_c \cdot \vec{x}) < 1 \\ 0 & \text{else} \end{cases}$ |
| MAE | $\vec{\delta}_c = \begin{cases} \alpha \vec{x} & \text{if } (\vec{w}_c \cdot \vec{x}) < r \\ -\alpha \vec{x} & \text{else} \end{cases}$ |
| Square | $\vec{\delta}_c = \alpha (r - \vec{w}_c \cdot \vec{x}) \vec{x}$ |
| Exponential | $\vec{\delta}_c = \alpha r \exp(-r \vec{w}_c \cdot \vec{x}) \vec{x}$ |
| Cross-entropy | $\vec{\delta}_c = \alpha r \left( \frac{4}{1 + \exp(r \vec{w}_c \cdot \vec{x})} \right) \vec{x}$ |
| REINFORCE | $\vec{\delta}_c = \alpha r \left( 1 - \exp \left( \vec{w}_c \cdot \vec{x} - \log \left( \sum_{j=1}^C \exp(\vec{w}_j \cdot \vec{x}) \right) \right) \right) \vec{x}$<br>$\vec{\delta}_{i \neq c} = -\alpha r \exp \left( \vec{w}_i \cdot \vec{x} - \log \left( \sum_{j=1}^C \exp(\vec{w}_j \cdot \vec{x}) \right) \right) \vec{x}$ |

Except for REINFORCE, each rule above only changes the weight vector for the selected choice *c*; the other weight vector(s) are left unchanged 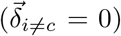. Additionally, the Exponential plasticity rule performs a weight normalization step (not shown): if the norm of the weights exceed a certain threshold (∥*w*_*i*_∥ *> B*), the weights are projected to the closest weight vector 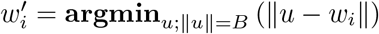 with 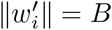. In this study, we chose *B* = 10.

Any remaining residual is assigned to *γ*_*ij*_ (i.e. corresponding to any interaction between rule and encoding stage). The importance of each model component could be assessed by calculating the proportion of variation in model scores that could be explained by the selection of component alone.

### S2 Appendix. Subtask consistency

In our primary benchmark, we measured human learning over 64 distinct subtasks, each consisting of 100 trials. For each subtask, the trial-averaged accuracy is a measure of the overall “difficulty” of learning that subtask, ranging from chance (0.5; no learning occurred over 100 trials) to perfect one-shot learning (0.995, perfect performance after a single example). For each of the 64 subtasks, one may estimate their trial-averaged performances (obtaining a length 64 “difficulty vector”), and use this as the basis of comparison between two learning systems (e.g. humans and a specific model).

To do so, we computed Spearman’s rank correlation coefficient (*ρ*) between a model’s difficulty vector and the human’s difficulty vector. The value of *ρ* may range between -1 and 1. If *ρ* = 1, the model has the same ranking of difficulty between the different subtasks (i.e., finds the same subtasks easy and hard). If *ρ* = 0, there is no correlation in the rankings.

In addition to computing *ρ* between each model and humans, we estimated the *ρ* that would be expected between two independent repetitions of the experiment we conducted here (i.e., an estimate of experimental reliability in measuring this difficulty vector). To do this, we took two independent bootstrap resamples of the experimental data, calculated their respective difficulty vectors, and computed the *ρ* between them. We repeated this process for *B* = 1, 000 bootstrap iterations, and thereby obtained the expected distribution of experimental-repeat *ρ*.

### S3 Appendix. Individual variability in overall learning ability

In this work, we focused primarily on *subject-averaged* measurements of human learning. However, individual subjects may also systematically differ from each other.

We aimed to investigate whether any such differences existed in learning behavior for the subtasks we tested in this study.

Here, we attempted to reject the null hypothesis that all subjects had the same learning behavior. To do so, we tested whether there were statistically significant differences in *overall learning performance* between individuals – that is, whether some individuals were “better” or “worse” learners. If this was the case, this implies individuals differ (at least in terms of overall learning performance), and the null hypothesis could be rejected.

### Permutation test for individual variability in overall learning ability

To test this null hypothesis, we identified a subset of human subjects who conducted all 64 subtasks in the primary, high-variation benchmark (n=22 subjects). For each subject, we computed their “overall learning performance”, which was their empirically observed average performance over all n=64 subtasks. That is, for subject *s*, we computed:

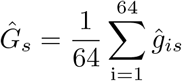

Where *ĝ*_*is*_ is the trial-averaged performance on subtask *i*, for subject *s*. The value of *Ĝ*_*s*_ is a gross measure of the subject’s ability to learn the objects in this study, ranging from 0.5 (no learning on all subtasks) to 0.995 (perfect one-shot learning on all subtasks). In total, we computed n=22 estimates of *Ĝ*_*s*_ (one for each subject in this analysis).

We then computed the sample variance over the various *Ĝ*_*s*_:

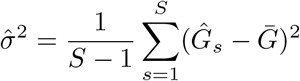

Where 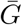 is the mean of overall lifetime performances. Intuitively, 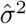is high if individuals differ in their overall learning performance, and is low if all individuals have the same overall learning performance (as would be the case under the null hypothesis).

We performed a permutation test on 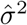 to test whether it was significantly higher than would be expected under the null hypothesis, permuting the assignments of each *ĝ*_*is*_ to each subject *s*. For each permutation, we computed the replication test statistic 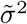 (using the same formulas above, on the permuted data). We performed *P* = 10, 000 permutation replications, then computed the one-sided achieved significance level by counting the number of replication test statistics greater than the actual, experimentally observed value 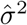.

### Testing whether specific humans outperform a model

To test whether a specific human has significantly higher overall learning abilities than a specific model (over the subtasks tested in this study), we performed Welch’s t-test for unequal variances on the overall learning performance, *Ĝ* (defined above). That is, for a specific subject *s* and model *m*, we attempted to reject the null hypothesis that *Ĝ*_*s*_ ≤ *Ĝ*_*m*_.

We adjusted for multiple comparisons using the Bonferroni correction (using the total number of pairwise comparisons we made between a model *m* and specific subjects *s*).

S1 Fig. In-lab vs. online behavioral measurements.

S2 Fig. Eyetracking measurements.

**Fig S1.**
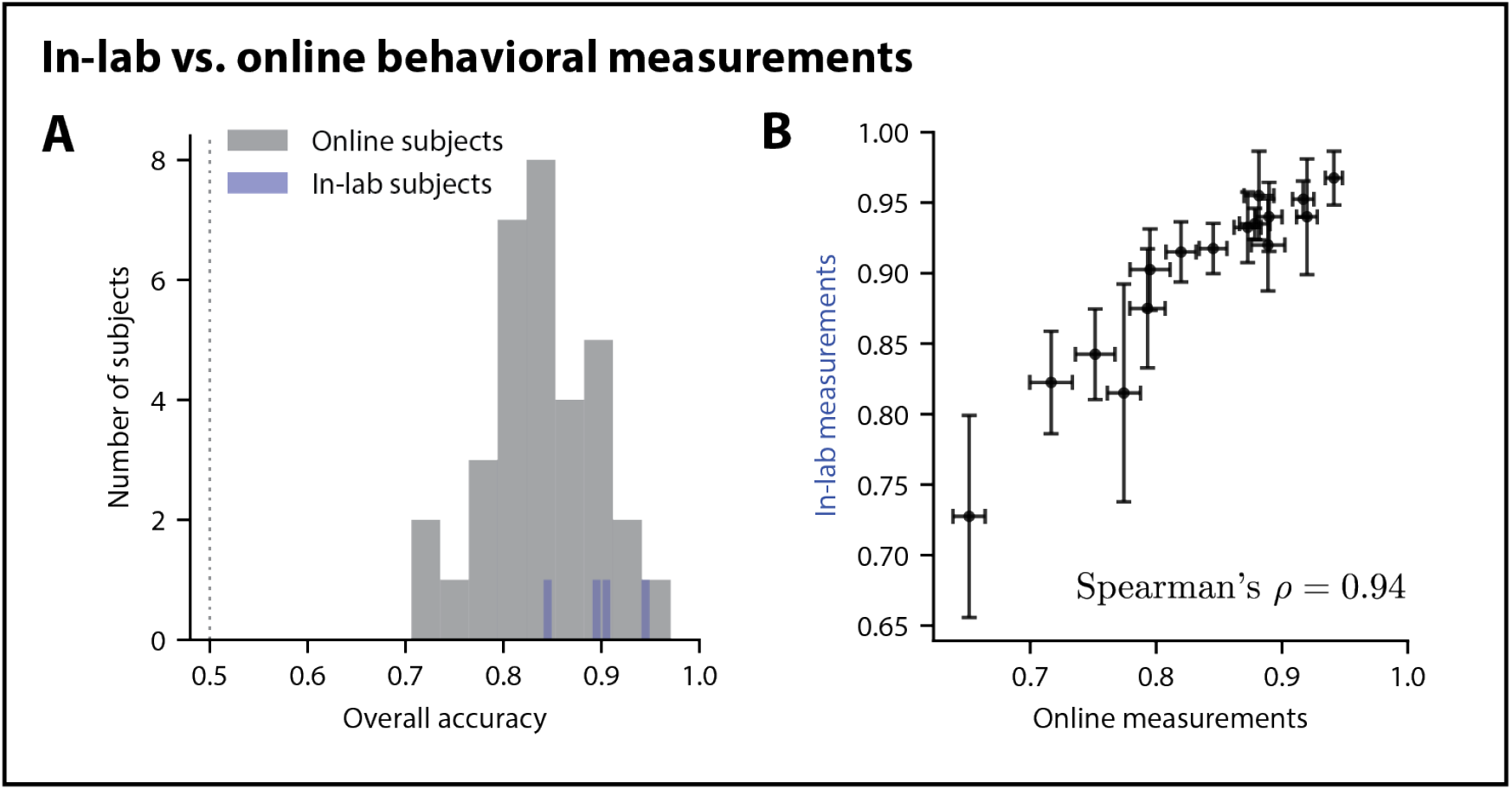
Comparison of online learning measurements with in-lab learning measurements. We took measurements for n=16 randomly selected subtasks from Experiment 1 in a group of in-lab human subjects (n=4) that used a chinrest and calibrated monitor setup. In **A**, we show that the overall accuracy of these in-lab subjects fell within the empirical support of the subject distribution from our online experiments. In **B**, we show that patterns of average accuracy (over subtasks) were tightly correlated between the in-lab and online populations (Spearman’s *ρ* = 0.94; see S2 Appendix). Errorbars are SEM (simple bootstrap over subjects).

**Fig S2.**
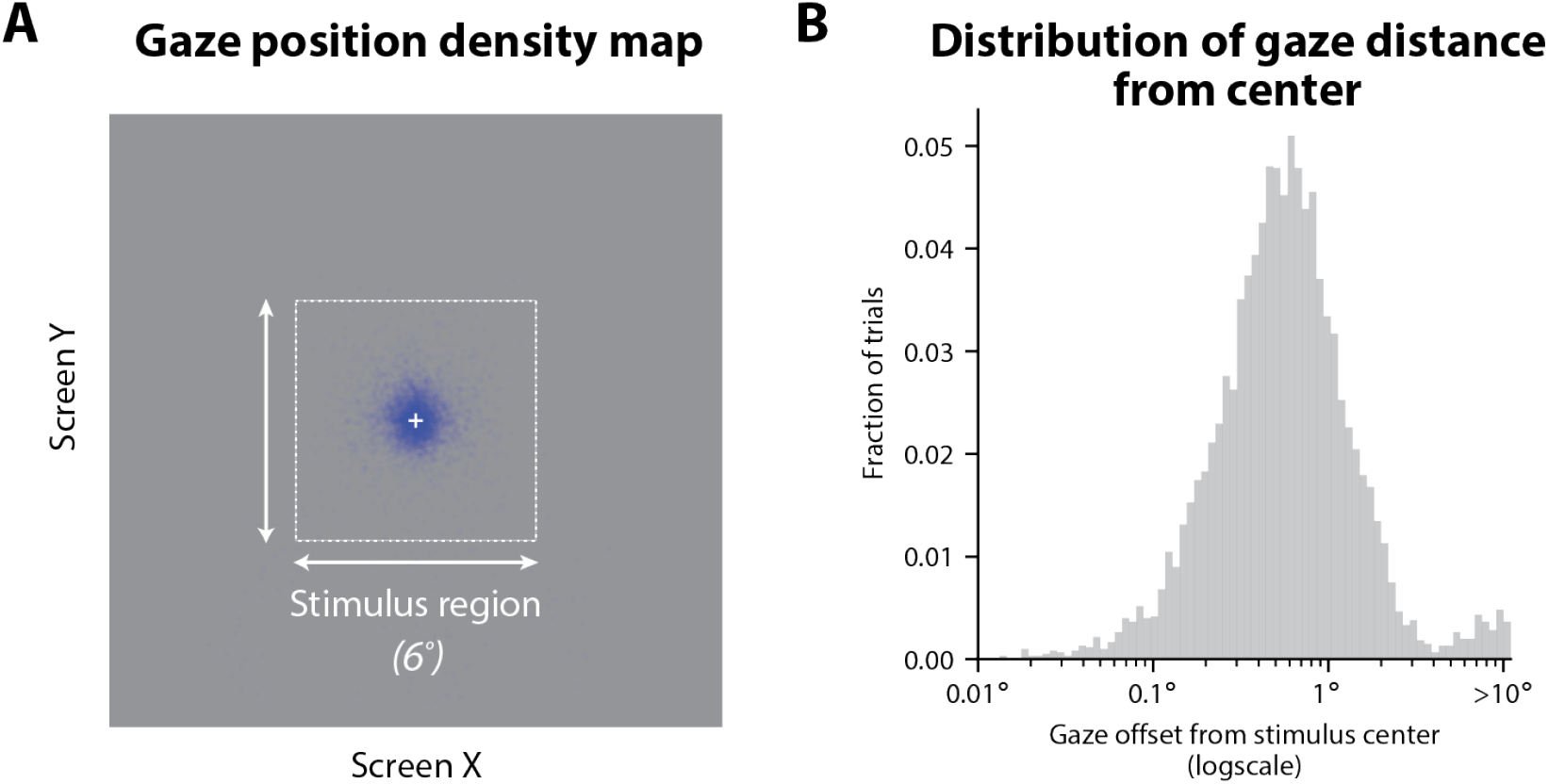
Distribution of gaze locations during learning task. We passively recorded eye movements from our in-lab subjects using an Eyelink 1000 Plus (monocular; desktop mounted) as they performed the task. In **A** is the overall distribution of the subjects’ gaze position (shown in blue) at the time of onset of stimulus presentation (i.e. distribution over subjects and trials). In **B** is the distribution of gaze distance from stimulus center (logscale) over all subjects and trials; the median distance from the center of the stimulus was 0.57° *±* 0.13° (mean *±* standard deviation over subjects). We found that on ≈95% of trials, the subject’s gaze was located in the test image region when it appeared on the screen.

## Footnotes

1 Except for models which are either at chance or perform perfect one-shot learning in all situations; then the correlation coefficient must be undefined.

2 The center of the test image is not necessarily the same as the center of the object in the test image.

3 We assumed our subjects used computer monitors with a 16:9 aspect ratio, and naturally positioned themselves so the horizontal extent of the monitor occupied between 40*°*-70*°* degrees of their visual field. Under that assumption, we estimate the visual angle of the stimulus would vary between a minimum and maximum of ≈ 4*°* − 8*°*. Given a monitor has a 60 Hz refresh rate, we expect the actual test image duration to vary between ≈ 183 − 217 milliseconds.

4 In practice, this was quite rare and corresponded to ≈0.04% of all trials that are included in the results in this work.

5 This can be seen by the expression for the variance of 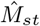, which is a mean over independent (but not necessarily identically distributed) Bernoulli variables: 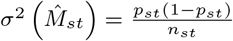. The value of 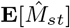 is the expected behavior of the model on trial *t* of subtask *s*, and *n*_*st*_ is the number of model simulations.

6 And/or because more model simulations were performed – though in this study, all tested models performed the same number of simulations, n=500.

7 In practice, this correction was relatively small, because of the high number of simulations that were conducted.

8 This estimator is biased.

9 The REINFORCE plasticity rule is a “policy gradient” rule that optimizes parameters directly against the rate of reward; it does not aim to predict reward.

